# Sordarin bound eEF2 unlocks spontaneous forward and reverse translocation on CrPV IRES

**DOI:** 10.1101/2023.03.04.530920

**Authors:** Zheren Ou, Alexey Petrov

## Abstract

The Intergenic Region Internal Ribosome Entry Sites (IGR IRESs) of *Discistroviridae* promote protein synthesis without initiation factors, with IRES translocation by elongation factor 2 (eEF2) being the first factor catalyzed reaction. Here, we developed a system that allows for the observation of intersubunit conformation of eukaryotic ribosomes at the single-molecule level. We use it to follow translation initiation and subsequent translocation of the cricket paralysis virus IRES (CrPV IRES). We observed that pre-translocation 80S-IRES ribosomes spontaneously exchanged between non-rotated and semi-rotated conformations but predominantly occupied a semi-rotated conformation. In the presence of eEF2, ribosomes underwent forward and reverse translocation. Both reactions were eEF2 concentration dependent, indicating that eEF2 promoted both forward and reverse translocation. The antifungal sordarin, stabilizes eEF2 on the ribosome after GTP hydrolysis in an extended conformation. 80S-CrPV IRES-eEF2-sordarin complexes underwent multiple rounds of forward and reverse translocations per eEF2 binding event. In the presence of sordarin, GTP hydrolysis or phosphate release were not required for IRES translocation. Together, these results suggest that in the presence of sordarin, eEF2 promotes the mid and late stages of CrPV IRES translocation by unlocking ribosomal movements, with mid and late stages of translocation being thermally driven.

## INTRODUCTION

Viruses employ alternative translation initiation mechanisms to translate their mRNAs. Internal ribosome entry sites (IRESs) are highly structured, untranslated regions of the viral mRNA that promote efficient cap-independent translation initiation (1, 2). The intergenic region (IGR) IRESs of Dicistroviruses are the simplest IRESs that start translation in the absence of all initiation factors (3–5). The most studied IGR IRESs are cricket paralysis virus (CrPV), taura syndrome virus (TSV), and plautia stali intestine virus (PSIV) IRESs that all initiate through a similar mechanism. It begins with IRES directly binding to the 40S ribosomal subunit, forming the 40S-IRES complex (4). This complex then recruits the large ribosomal subunit. In the resulting 80S-IRES ribosomes, the IRES occupies the intersubunit space spanning from the A to E site where it interacts with the phylogenetically-conserved 80S core (6–8). Consequently, CrPV IRES is being active in a broad range of organisms and can promote translation in insect cells (9, 10), mammalian cells and extracts (4, 11), and yeasts (12, 13). It is also capable of recruiting bacterial ribosomes. However, the reading frame is not established in bacteria and translation begins downstream from the AUG codon (14). CrPV IRES is composed of three structural domains (15). Domains I and II recruit the ribosome (11, 16). Domain III, composed of RNA pseudoknot I (PKI) with the GCU translation start codon, is located immediately downstream. In 80S-IRES complexes, PKI occupies the A-site of the ribosome and spans into the decoding center, where it mimics the anticodon stem of tRNA base paired to a mRNA (17, 18). Thus subunit joining results in pre-translocation ribosomes and, 80S-IRES complex must be translocated to allow the first codon decoding (18–21). IRES translocation requires eEF2 and places the first GCU codon into the A site, where it is decoded by eEF1A-tRNA^Ala^.

Translocation is accompanied by large conformational changes of the ribosome. Upon peptide bond formation, acceptor ends of the A- and P-site tRNAs move into hybrid A/P and P/E states (22, 23). Transition into the hybrid state is concurrent with an ∼9 degrees counterclockwise (forward) rotation of the small subunit relative to the large subunit (23, 24). Simultaneously, the L1 stalk moves inward and contacts the elbow of the P-site tRNA (25–27). The pre-translocation ribosomes spontaneously exchange between the classical and hybrid states (28, 29), with the hybrid state being favored at physiological magnesium and polyamine concentrations (30, 31). EF-G binding triggers conformational rearrangements of the ribosome (32), with the ribosomes transiently occupying the rotated state prior to GTP hydrolysis (26, 28, 33, 34). In the rotated ribosomes, domain IV of EF-G extends into the A-site and the head of the 30S subunit swiveled by 18 degrees. These rearrangements are accompanied by movement of the anticodon stem-loop, placing tRNA into chimeric ap/P and pe/E sites (35–37). Post-GTP hydrolysis, the body of the small subunit undergoes reverse (clockwise) rotation and the head of the small subunit swivels back. The combination of these two motions leads to translocation of tRNAs into the P- and E-sites (35, 38). During translocation, the L1 stalk maintains contact with the tRNA (26, 39). In bacteria and higher eukaryotes, after translocation is complete, the L1 stalk moves outward, allowing the E-site tRNA to dissociate (26, 40, 41). However, in fungi, after tRNA movement is complete, the L1 stalk remains in a closed position and its opening is facilitated by the elongation factor 3 (eEF3) (42, 43).

The translocation of 80S-IRES complex bears similarity to regular tRNA translocation. The 80S-CrPV IRES pre-translocation complexes occupy non-rotated and semi-rotated conformations that differ by 5 degrees counterclockwise rotation of 40S subunit body (18, 19). It was proposed that these conformations are at equilibrium that resemble the semi-rotated - non-rotated state exchange in pre-translocation ribosomes. Reminiscent to tRNA translocation, eEF2 binding to 80S-CrPV IRES complexes induces an additional 3 degrees counterclockwise rotation, resulting in fully rotated ribosomes. Simultaneously, tRNA-mRNA mimicking PKI moves into the chimeric ap position on the small subunit (17). The Cryo-EM structure of 80S ribosomes with related TSV IRES (44, 45), eEF2, and the antifungal sordarin, showed ribosomes in five intermediate stages of IRES translocation (46). There, the reverse body rotation and forward head swivel are accompanied by progressive movements of PKI toward the P-site. This movement is followed by reverse head swivel that coincides with final placement of the PKI in the P-site, yielding the translocated IRES and ribosome in the non-rotated state (17, 46). The translocated 80S-CrPV IRES complex is not stable and back translocates, as was shown by both toeprinting analysis and Cryo-EM (5, 47, 48). Despite these studies of 80S-CrPV IRES complex, the molecular mechanism of IRES initiation and translocation remains unclear. This is attributed in part to the lack of ability to track the conformation of eukaryotic ribosomes in real-time.

Here we report the development and application of a single-molecule FRET system that allows us to observe intersubunit conformation of eukaryotic ribosomes in real-time. We used it to follow translation initiation and translocation of CrPV IRES. We showed that 60S subunit joining places ribosomes in both semi-rotated and non-rotated conformations. Pre-translocation 80S-CrPV IRES complexes are exchanging between these two conformations. Using ribosomal rotation as a proxy for translocation, we showed that both forward and reverse translocation are driven by eEF2. GTP hydrolysis is required for rapid forward translocation. However, in the presence of the antifungal sordarin, that stabilizes eEF2 on the ribosome, GTP hydrolysis or phosphate release were not needed, and forward and reverse translocation occurred spontaneously and repeatedly at rates that were comparable with normal translocation. Thus, sordarin captures ribosome in an unlocked state in which the mid and late steps of IRES translocation are thermally driven. The experiments with GDP and GDPNP indicated that GTP hydrolysis is likely needed to achieve an unlocked state. Together these results provide support for the Brownian model of translocation.

## MATERIAL AND METHODS

### rRNA Mutagenesis, mutant selection, and validation

Ribosome mutagenesis was performed, as previous described (49), using the RDN mutagenesis system (50, 51). Helix 44 of 18S rRNA was extended, as described before, and amplified from the plasmid pAL7 (a plasmid contains h44 mutant) with F_h44_SexAI and R_h44_MluI primers (Supplementary Table 1). The PCR product was cloned into the RDN operon bearing pJD180. Trp plasmid (52) by SexAI and MluI sites. Mutations on helix 101 of 25S rRNA were introduced into the plasmid pJD180.Trp by two-step megaprimer PCR. First, primers F_h101_b and R_h101.3_b were used to amplify the downstream helix101 mutant, primers F_h101.3_a and R_h101_a were used to amplify the upstream helix 101 mutant. Upstream and downstream PCR products were annealed together to get the full helix 101 mutant insertion. The insertion bearing hairpin sp22 sequence was cloned into pJD180. Trp by BamHI and MluI sites. The resulting plasmids pAL783 (pJD180.Trp.h44.sp68) and pAL797 (pJD180.Trp.h101.3.sp22) were transformed into the S. cerevisiae strain AL14 (49), which expressed mutant rRNA from pJD694 (URA3). Transformants were initially selected on −Trp medium, followed by two passages of replica plating onto 5-FOA medium. RDN mutagenesis and lack of wild type loci were confirmed by RDN PCR and sequencing. A pair of primers, which are about 150 bp up and downstream of the insertion site, were used to amplify the insertion region. The product is 383 bp for the helix 44 mutant and 336 bp for the helix 101 mutant, which are 19 bp and 21 bp longer than the PCR product from the wildtype RDN operon. The PCR product was analyzed by electrophoresis on 4% agarose gel, which migrated as a single band with a size corresponding to the mutant hairpin. Furthermore, sequencing of the PCR product confirmed the presence of the mutant hairpin and absence of the wildtype contamination.

### Purification and labeling of ribosomal subunits

Ribosomal subunits were purified and labeled, as previously described (49, 53), with the following modifications: A 5 ml liquid YPAD culture was inoculated with a single colony from a fresh YPAD plate and grown at 250 rpm in a 30°C shaker until OD600 = 0.8-1. 1 ml of the resulting culture was used to inoculate 1 L of YPAD media. Cells were grown until OD600 = 0.8-1.0, harvested by centrifugation, and washed twice with ice-cold buffer A (30 mM HEPES-KOH pH = 7.4, 100 mM KCl, 15 mM MgCl_2_, 2 mM DTT). Cells were lysed by a bead-beater and the lysate was cleared by centrifugating twice in an A27-8×50 rotor for 15 min at 15,000 rpm. Fines were removed by passing the lysate through a 0.45 µm filter. The lysate was overlayed on a 3 ml cushion of buffer B (30 mM HEPES-KOH pH = 7.4, 100 mM KCl, 12 mM MgCl_2_, 2 mM DTT), supplemented with 1 M sucrose and spun down in a Type 70Ti rotor for 5 h at 55,000 rpm. The ribosome pellet was resuspended in buffer B and loaded on top of 5%-47% sucrose gradients in Buffer B. Gradients were centrifuged at 22,000 rpm for 12 h in a SW32 rotor. Gradients were fractionated and 80S ribosomes were collected and buffer exchanged into 80S buffer B supplemented with 250 mM sucrose. 80S ribosomes were aliquoted and frozen with liquid nitrogen. Purified 80S ribosomes were labeled with corresponding oligonucleotide dye at a 1:2 molar ratio by sequentially incubating at 42°C for 1 min, 37°C for 10 min, and 30°C for 10 min. The reaction was loaded on top of 10%-30% sucrose gradients (30 mM HEPES-KOH pH = 7.4, 500 mM KCl, 7.5 mM MgCl_2_, 2 mM DTT supplement with sucrose) and centrifugated at 24,000 rpm for 12 h in a SW32 rotor. Gradients were fractionated and 40S and 60S ribosomal subunits were collected. Labeled ribosomal subunits were concentrated and buffer exchanged into buffer D (30 mM HEPES-KOH pH = 7.4, 100 mM KCl, 5 mM MgCl_2_, 2 mM DTT, and 250 mM sucrose) using Amicon® Ultra-15 100 KDa centrifugal filters. Ribosomal subunits were aliquoted and frozen with liquid nitrogen. The purity and integrity of subunits was validated by composite agarose-acrylamide gel electrophoresis. The labeling efficiency was measured spectrophotometrically and was found to be 92% and 88% for 40S and 60S subunits, correspondingly. The measurement was confirmed by quantifying the fluorescence intensity of the 40S-Cy3B and 60S-Cy5 subunit bands of the composite gel.

### Purification of elongation factors

eEF1A, eEF1Bα, eEF2, and eEF3 were purified, as previously described (53–56).

### eEF1A

Untagged eEF1A was purified from the yeast strain, CB010, as previous described (54) with the following modifications: The lysate was clarified by centrifugation in an A27-8×50 rotor at 15,000 rpm for 30 min, followed by centrifugation in a Type 70 Ti rotor at 50,000 rpm for 90 min. The clarified lysate was applied to a gravity-flow 7 ml DEAE column equilibrated with eEF1A buffer A (20 mM Tris-HCl, pH 7.5, 100 mM KCl, 0.1 mM EDTA, 1 mM DTT, 25% glycerol). After loading the lysate, the column was washed with two column volumes (CV) of eEF1A buffer A. The flow-through and column wash were pooled and loaded to a gravity-flow 5 ml SP Sepharose column equilibrated with eEF1A buffer A. The SP Sepharose column was washed with 5 CV of eEF1A buffer A before eluting. eEF1A was eluted with eEF1A buffer 150B (eEF1A buffer A with 150 mM KCl) for 5CV at first, then eluted with buffers 250B, 350B, and 500B for 4 CV. 5ml fractions were collected. eEF1A containing fractions were pooled and were concentrated to ∼5 ml. The concentrated protein was applied to a 26/600 Superdex 75 size exclusion column equilibrated with a storage buffer (20 mM Tris-HCl, pH 7.5, 100 mM KCl, 0.1 mM EDTA, 1 mM DTT, 25% glycerol). Fractions containing pure eEF1A were identified by SDS-PAGE, pooled, and concentrated using Amicon® Ultra-15 30 kDa MWCO centrifugal filter units.

### eEF1Bα

pET28b plasmid carrying eEF1Bα-TEV site-His6 was transformed into BL21-RIPL *e.coli* cells. Protein was expressed and purified, as previous described (53).

### eEF2

6 x His tagged eEF2 was purified from the yeast strain, TKY 675, as previous described (55) with the following modifications. Cells were grown in YPAD at 30°C to OD600 = 1.5 and harvested by centrifugation. The cells were resuspended with ice-cold lysis buffer (50 mM potassium phosphate, pH = 7.6, 300 mM KCl, 1 mM DTT, 10 mM imidazole) and lysed using a bead-beater. After lysis, the pH of the lysate was adjusted to 7.7 with 1 M un-titrated Tris and clarified by centrifugation in an A27-8×50 rotor at 12,500 rpm for 20 min. The resultant supernatant was centrifuged at 50,000 rpm in a Type 70 Ti rotor for 90 min. The lysate was passed through a 0.22 µm filter and loaded onto a 5 ml HisTrap HP column (GE Healthcare). The column was washed with 25 ml of lysis buffer containing 30 mM imidazole. Proteins were eluted with lysis buffer containing 250 mM imidazole. Elution was exchanged into Buffer A (20 mM Tris-HCl pH 7.6, 30 mM KCl, 5 mM MgCl_2_, 1 mM DTT). Protein was loaded on to a 5 ml HiTrap Q column. Column was washed with 6 CV of buffer A. Proteins were eluted with a 15 CV gradient of 30-500 mM KCl. Fractions containing eEF2 were pooled and concentrated using Amicon® Ultra-15 30 kDa MWCO centrifugal filter and exchanged into storage buffer (20 mM Tris-HCl, pH 7.5, 100 mM KCl, 0.1mM EDTA, 1 mM DTT, 25% glycerol).

### eEF3

6 x His tagged eEF3 was purified from the yeast strain, TKY 1653, as previous described (56).

### Preparation and biotinylation of CrPV IRES

CrPV IRES was prepared by T7 runoff transcription. The template plasmid T7-IGR-CrPV-IRES, was linearized with NarI and transcribed with NEB HiScribe™ T7 High Yield RNA Synthesis Kit. The resulted RNA was purified by gel filtration on a SEC650 column, as described before (57). RNA was concentrated using a 30 kDa MWCO spin concentrator. The 5′-biotinylated DNA oligonucleotide was annealed to the 3′ end of the RNA to the region 39-54 nts downstream of the PKI of CrPV IRES, as described before (53).

### Single-molecule imaging and data analysis

A home-built total internal refection fluorescence (TIRF) microscope was used for single-molecule imaging. PEG-Biotin treated quartz slides were washed twice with 200 μl of TP50 buffer (50 mM Tris-HCl, pH 7.5, 100 mM KCl) and incubated with 1 μM neutravidin, 0.67 mg/ml BSA, and 1.3 μM blocking DNA oligonucleotide duplex for 5 min. Slides were washed twice with 200 µl of TP50 and one time with a reaction buffer containing 30 mM HEPES-KOH, pH = 7.4, 100 mM KCl, 5 mM MgCl_2_, and 1 mM Spermidine. 40S-CrPV IRES complex were prepared by incubating 100 nM of 40S-Cy3B fluorescent subunits and 100 nM CrPV IRES-biotin in a reaction buffer at 30°C for 5 min. 40S-CrPV IRES complexes were diluted to 0.8 nM with reaction buffer and immobilized for 5 min. Non-immobilized ribosomes were removed by washing the chambers with reaction buffer. Finally, the buffer was exchanged to reaction buffer supplemented with 1mM GTP, 2 mM TSY, and an oxygen scavenging system composed of 2.5 mM protocatechuic acid and 0.06 U/μl protocatechuate dehydrogenase (58). A 50 μl delivery mix was flowed to the slide simultaneously with the start of observation. For experiments with sordarin (Cayman Chemical #26255), drug concentration was 20 μM. All experiments were performed at 21°C.

Fluorescent traces were extracted using home-written Matlab scripts. Time-dependent donor and acceptor fluorescence intensities were converted into FRET efficiency using the formula *FRET* = *I_acceptor_*/(*I*_*acceptor*_ + *I*_*donor*_). Single molecules were identified by single step photobleaching events. FRET trajectories were idealized using ebFRET software (59, 60). According to ELBO criterion, the three-state model (two FRET states and FRET off) was preferable in the majority of experiments. However, the minority of traces (∼1%) clearly showed three distinct FRET states. signal-to-noise hampered identification of this third state in in majority of the molecules. Therefore, to avoid overinterpretation of the results, we selected a simpler two FRET state model.

All statistical analysis was performed in Matlab. Kinetic parameters were extracted by building cumulative probability plot and fitting observed data to linear, single- or double-exponential functions. The reported 95% confidence interval is a goodness of a fit. The rotating molecules were defined as molecules with at least two unambiguous FRET transitions between the fitted FRET states. All related data size and statistical parameters were reported in Supplementary Table 2-5.

## RESULTS

### Following 80S-CrPV IRES complex formation in real time

To follow conformation of eukaryotic ribosomes in real-time, we site-specifically labeled yeast small and large ribosomal subunits with fluorescent dyes. Phylogenetically variable loops of helix 44 of 18S rRNA and helix 101 of 25S rRNA were extended with distinct metastable hairpins (Figure 1A). Isolated strains were confirmed to have only mutant rDNA by PCR and RDN loci sequencing (Supplementary Figure 1). These hairpins allow site-specific annealing of fluorescent DNA oligonucleotides (49, 61). Small ribosomal subunits were site-specifically labeled with Cy3B dye and large ribosomal subunits were labeled with Cy5 dye by annealing fluorescently labeled DNA oligonucleotides to the h44 and h101 extensions correspondingly (Supplementary Figure 1).

**Figure 1.**
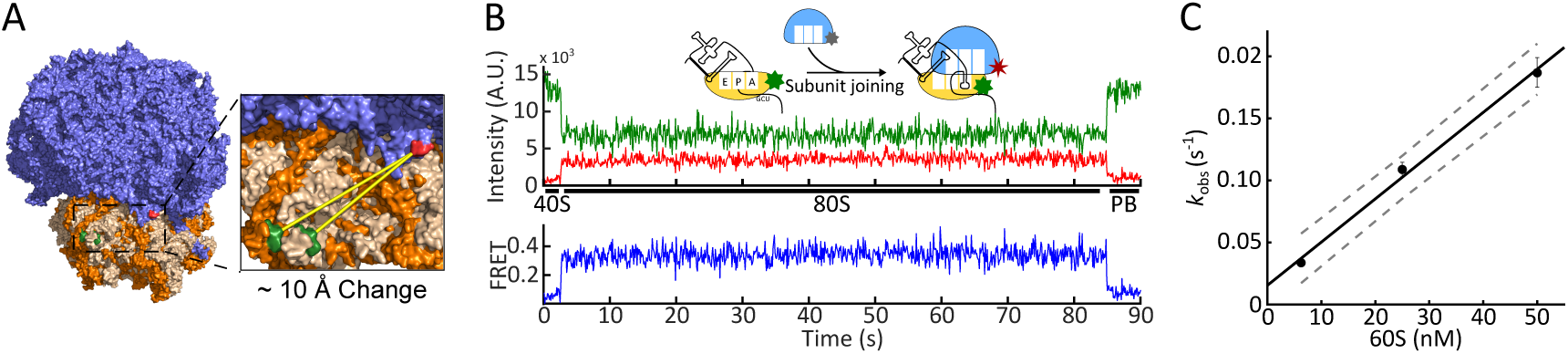
Following 80S-CrPV IRES formation in real time. A. 40S (Orange) and 60S (Blue) yeast ribosomal subunits were labeled at h44 (Green) and h101 (Red), correspondingly. The distance between dye positions is expected to change by ∼10 Å when the ribosome transits between the rotated and non-rotated conformations. Based on PDB IDs 3J77 and 3J78 (62). B. FRET between h44 and h101 attached dyes was used to follow 60S subunit joining to 40S-IRES complexes. The 40S-IRES complexes were immobilized on the slide and 60S subunits were delivered concurrently with the start of observation. 80S ribosome formation was detected by an appearance of FRET (∼3 seconds in the example trace). FRET remained stable as framerate is insufficient to detect spontaneous rotations. C. 60S subunit arrival rate was concentration dependent. The arrival rate constant was measured as a slope of the concentration dependence curve and was determined to be 3.5 ± 1.0 µM^-1^s^-1^. Error bars are the errors of the exponential fits (n = 102, 130, and 117 molecules for 6.25 nM, 25 nM, and 50 nM 60S ribosomal subunits).

The Cryo-EM structures of both 80S ribosomes in complex with tRNAs and IGR IRESs show that apical positions of h44 and h101 are within FRET distance and are ∼60Å apart in non-rotated conformation (19, 46, 62). The distance between dyes increases by ∼10 Å when the ribosome transitions from non-rotated to rotated conformation (PDB IDs 3J77 and 3J78) (62). The increase in distance will result in lower FRET efficiency. This allows us to follow 60S subunit joining by appearance of FRET and ribosome conformation by measuring FRET efficiency.

Using this system, we first followed 80S formation on CrPV IRES. The 40S-CrPV IRES complex is exceedingly stable with a lifetime of more than 400 s (53). This allowed us to prepare, and surface immobilize 40S-Cy3B-CrPV IRES complexes using biotinylated CrPV IRES. Then, Cy5-labeled large ribosomal subunits were delivered simultaneously with the start of observations and 80S ribosome formation was detected by an appearance of FRET (Figure 1B). 60S ribosomal subunits binding was efficient, with 70 % of the 40S-IRES ribosomes forming 80S-IRES complexes. Large subunit arrival rate was fast with *k*_obs_ = 0.19 ± 0.012 s^-1^ at 50 nM 60S subunits (Supplementary Table 2). Consistent with biomolecular reaction, the arrival rate was concentration dependent (Supplementary Figure 2A). The rate constant was determined as a slope of concentration dependence and was found to be 3.5 ± 1.0 µM^-1^s^-1^ (Figure 1C). The measured rate is comparable with 60S recruitment rates during factor driven initiation measured by ensemble and single-molecule approaches (0.076 s^-1^ and 0.2 s^-1^, correspondingly, at 100 nM 60S ribosomal subunits (63, 64)). Consistent with previous studies (53), 80S-CrPV IRES complexes were stable, with lifetimes longer than the five minutes observation window (Supplementary Figure 2B). Thus, labeled ribosomal subunits are active in IRES driven initiation.

### Spontaneous rotations in pre-translocation 80S-CrPV IRES complex

The Cryo-EM structures show 80S-IRES complexes in both semi-rotated and non-rotated conformations. It was proposed that 80S-IRES complexes are spontaneously exchanging between these conformations (18, 19). In experiments described above, ribosomes predominantly occupied the low ∼ 0.22 FRET state and transiently transitioned into the higher ∼ 0.34 FRET state (Figure 2B). Cryo-EM structures of 80S IRES complexes showed that the distance between the end of helix 44 and the end of helix 101 decreases by ∼ 5 Å as the ribosome changes from semi-rotated to the non-rotated conformation (Supplementary Figure 3) (18, 19). Thus, we assigned the low (0.22) FRET state to the semi-rotated ribosomes and the high (0.34) FRET state to the non-rotated ribosomes. Consistently with two conformations, total FRET efficiency histogram of 80S-IRES complexes had a right shoulder and was best approximated by double Gaussian fit. The FRET states of double Gaussian fit matched individual FRET distributions and had efficiency of 0.2 ± 0.06 (95% CI) for the low FRET state and 0.33 ± 0.07 (95% CI) for the high FRET state (Supplementary Figure 2C).

**Figure 2.**
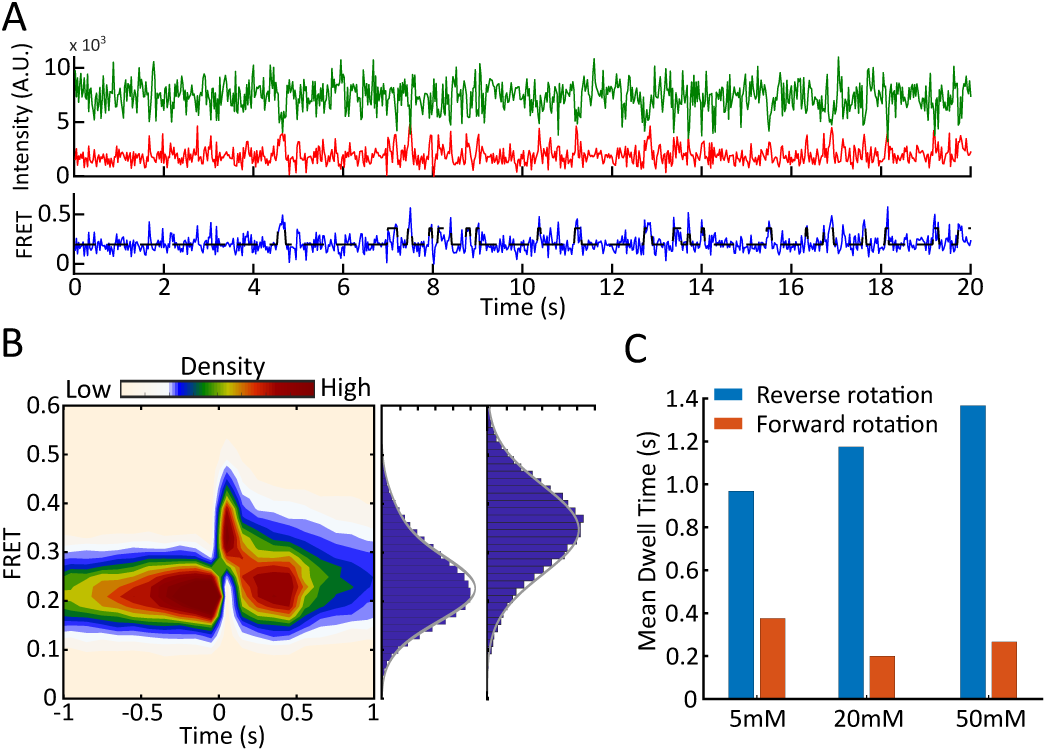
Spontaneous rotation of pre-translocation 80S-CrPV IRES complexes. A. Example trace showing spontaneous rotations of 80S-CrPV IRES complex. B. Density heatmap. FRET traces were post-synchronized to the point where semi-rotated ribosome transited to non-rotated ribosome set as time 0. The corresponding FRET distribution for semi-rotated ribosome and non-rotated ribosome were best described by a single Gaussian. The estimated mean FRET efficiency of 0.22 ± 0.08 (95% CI) for semi-rotated state and 0.34 ± 0.08 (95% CI) for high FRET state correspondingly. C. Bar chart of mean dwell times for reverse (blue) and forward (orange) rotations at various Mg^2+^ concentrations at 25ms exposure time.

Experiments described above were done at 100 ms exposure time. At this condition, 27% of 80S ribosomes showed spontaneous rotations, with an average of 2.3 rotations per trace (average FRET duration is 169 ± 10.3 s) (Supplementary Table 2). This is substantially less frequent than what was observed in pre-translocation bacterial and human ribosomes (28, 65). Importantly, spontaneous transitions of pre-translocation ribosomes in complex with tRNAs are fast and occur on the sub-second time scale (28, 29, 66–68). We hypothesized that spontaneous transitions (and particularly transient high state) are faster than 100 ms, and, thus, escaping detection due to time-averaging by the imaging camera (69). To determine if the low frame rate masked spontaneous transitions, we imaged 80S-CrPV IRES complex at 25 ms resolution. The increased time resolution made subunits delivery experiments challenging due to lower signal to noise ratio (SNR) and further decrease of the SNR due to mechanical stresses produced by the solvent delivery system. Therefore, 25 ms experiments were done with pre-formed 80S-CrPV IRES complexes. There, 83% of pre-translocation ribosomes underwent spontaneous rotations with an average of 23.3 rotations per molecule (Supplementary Table 2). The dwell times were best described by double exponential function (Supplementary Figure 4), indicating kinetic heterogeneity. Ribosomes highly preferred the semi-rotated state, with a mean dwell time being 0.97 s, while non-rotated state was approximately three times shorter at 0.38 s (Figure 2C). The median high FRET state dwell time was 0.05 s, which explains why spontaneous rotations were rarely observed at 100 ms exposure time and confirms that they have been masked by low framerate (Supplementary Table 2). Together this confirms the hypothesis that 80S-IRES complexes spontaneously exchange between semi-rotated and non-rotated conformations.

### eEF2 facilitates intersubunit dynamics

Translocation is accompanied by reverse subunit rotation and results in non-rotated ribosomes. Translocation of 80S-tRNA complexes is unidirectional and results in stable translocated ribosomes. However, translocated IRES complexes are unstable and back translocate (18, 48). To follow translocation of CrPV IRES, we co-delivered 60S subunits, eEF2, and GTP to the immobilized 40S-CrPV IRES complexes. Spontaneous rotations complicate identifications of conformational transitions associated with translocation. Thus, we conducted these experiments at 100 ms time resolution, where spontaneous transitions in pre-translocation complexes are masked by the camera averaging, and predominantly eEF2 induced rotations will be visible.

In the presence of eEF2-GTP, the 80S ribosomes began exchanging between the semi-rotated (0.23) and non-rotated (0.34) FRET states (Figure 3A) at rates that are drastically different from spontaneous rotations (Figure 2C, Supplementary Figure 5A). The semi-rotated state mean dwell time was 8.4 s at 100 nM eEF2 and 0.97 s in the absence of eEF2. The non-rotated state mean dwell time was 5.9 s at 100 nM eEF2 and 0.38 s in the absence of eEF2, indicating that these transitions were eEF2 induced (Supplementary Table 2, 3). Translocation places ribosome into the non-rotated, high FRET state. Thus, low to high FRET transition in the presence of eEF2 corresponds to the forward translocation. By extension, the high to low FRET transition corresponds to counterclockwise intersubunit rotation and thus back translocation.

**Figure 3.**
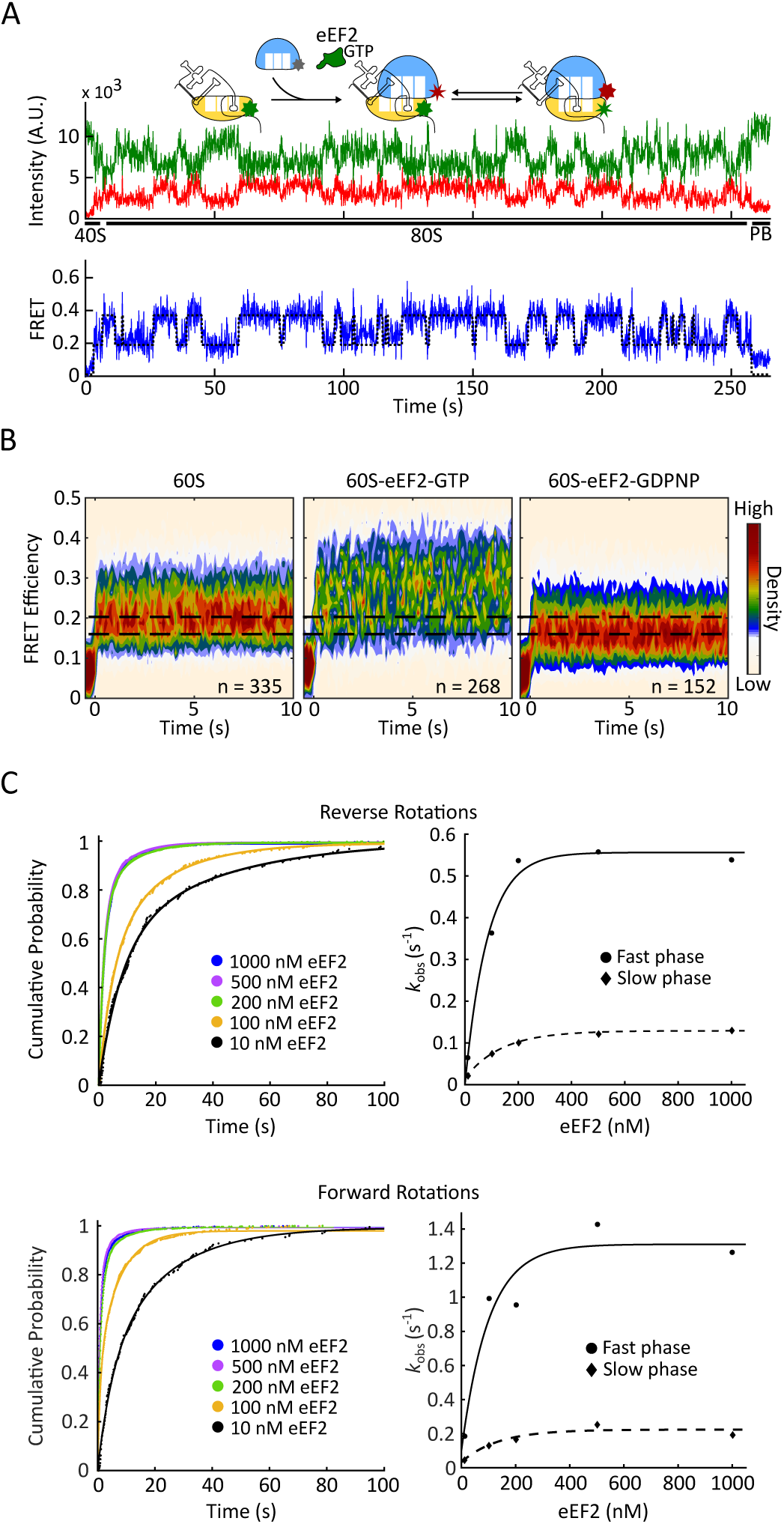
eEF2 facilitates intersubunit rotations. A. Example trace of 80S-CrPV assembly in the presence of eEF2-GTP. 40S-IRES complex were immobilized to the slide and 25 nM 60S subunits, eEF2 at various concentrations, and 1mM GTP were co-delivered concurrently at the start of observation. Upon subunit joining, FRET oscillated between high and low FRET states that correspond to non-rotated and semi-rotated ribosomal conformations. B. Density heatmap of FRET traces. FRET traces were post-synchronized to the beginning of FRET (60S subunit binding), with synchronization point being set to time 0. In the presence of GTP, ribosomes fluctuated between semi-rotated and non-rotated conformations. Ribosomes rapidly shifted into rotated state in the presence of GDPNP. C. Cumulative probability plot of reverse (top) and forward rotation (bottom) dwell times with different eEF2-GTP concentrations. Rotation rates saturated at around 200 nM eEF2. Lines represent double-exponential fitting, and the resulted rates were fit by hyperbolic functions (line, fast phase; dashed, slow phase). (n = 143, 129, 268, 202, and 169, correspondingly).

To understand the translocation mechanism, we varied eEF2 concentration in the delivery mix from 10 to 1000 nM. The rates of both forward translocation (low to high FRET) and surprisingly reverse (high to low FRET) translocation were dependent on eEF2 concentration indicating biomolecular mechanism. The kinetics of both reactions were remarkably similar. Both were best approximated by double exponential fits. The rates of both fast and slow components increased with eEF2 concentrations, and relative amplitude increased for the fast component and decreased for the slow component (Supplementary Table 3). Both fast and slow components saturated at ∼ 200 nM eEF2 for the forward and reverse rotations (Figure 3C). Interestingly, this is comparable with previously measured the K_m_ of GTP hydrolysis by EF-G (80 nM (70) to 400 nM (71)). Thus, eEF2 efficiently engages both conformations of 80S-IRES complex and promotes both forward and reverse translocation.

As a control, we replaced GTP with GDP and non-hydrolyzable GTP analog GDPNP. eEF2-GDPNP stabilizes ribosomes in fully rotated state that is characterized by additional 3° counterclockwise rotation (17) that should further decrease FRET efficiency. In agreement, 80S-IRES-eEF2-GDPNP complexes had a lower 0.17 ± 0.04 FRET (Figure 3B, Supplementary Table 5) that corresponds to the fully rotated ribosomes. Translocation in the presence of GDP is slow due to low affinity of eEF-2-GDP to the ribosome and absence of GTP hydrolysis. Consistent with this, eEF2-GDP had little effect on ribosome conformation. (Supplementary Figure 6A).

### Sordarin bound eEF2 accelerates intersubunit rotation

Sordarin is a tetracyclic diterpene glycoside that inhibits translation by stabilizing eEF2 on the ribosome (72). It does not inhibit GTP hydrolysis nor affects phosphate release (46, 72). It binds to the lever arm of eEF2 in the cleft between domains III, IV, and V and restricts compaction of eEF2, thus stabilizing eEF2 in the extended conformation and preventing eEF2 dissociation (73, 74). We used sordarin to probe the role of eEF2 in IRES translocation. First, we followed 80S formation in the presence of sordarin by delivering 60S ribosomal subunits and sordarin (no eEF2) to 40S-CrPV IRES complex. As expected, sordarin alone did not affect 80S complex formation and spontaneous rotations in pre-translocation 80S-IRES complexes (Supplementary Table 5). Then, we repeated these experiments with sordarin and eEF2. Similar to the experiments without the drug, eEF2-soradarin promotes intersubunit rotations, i.e. forward and reverse translocation (Figure 4A). However, reaction rates were greatly increased at low eEF2 concentrations. Furthermore, saturation concertation of eEF2 was 20 times lower in the presence of sordarin than in absence of the drug (10 nM vs 200 nM eEF2) (Figures 4C, Supplementary Table 4). Together, with previous reports showing that sordarin stabilizes eEF2 on the ribosome, this suggests that when sordarin is present, eEF2 promotes multiple rounds of forward and reverse rotations without dissociating from the ribosome.

**Figure 4.**
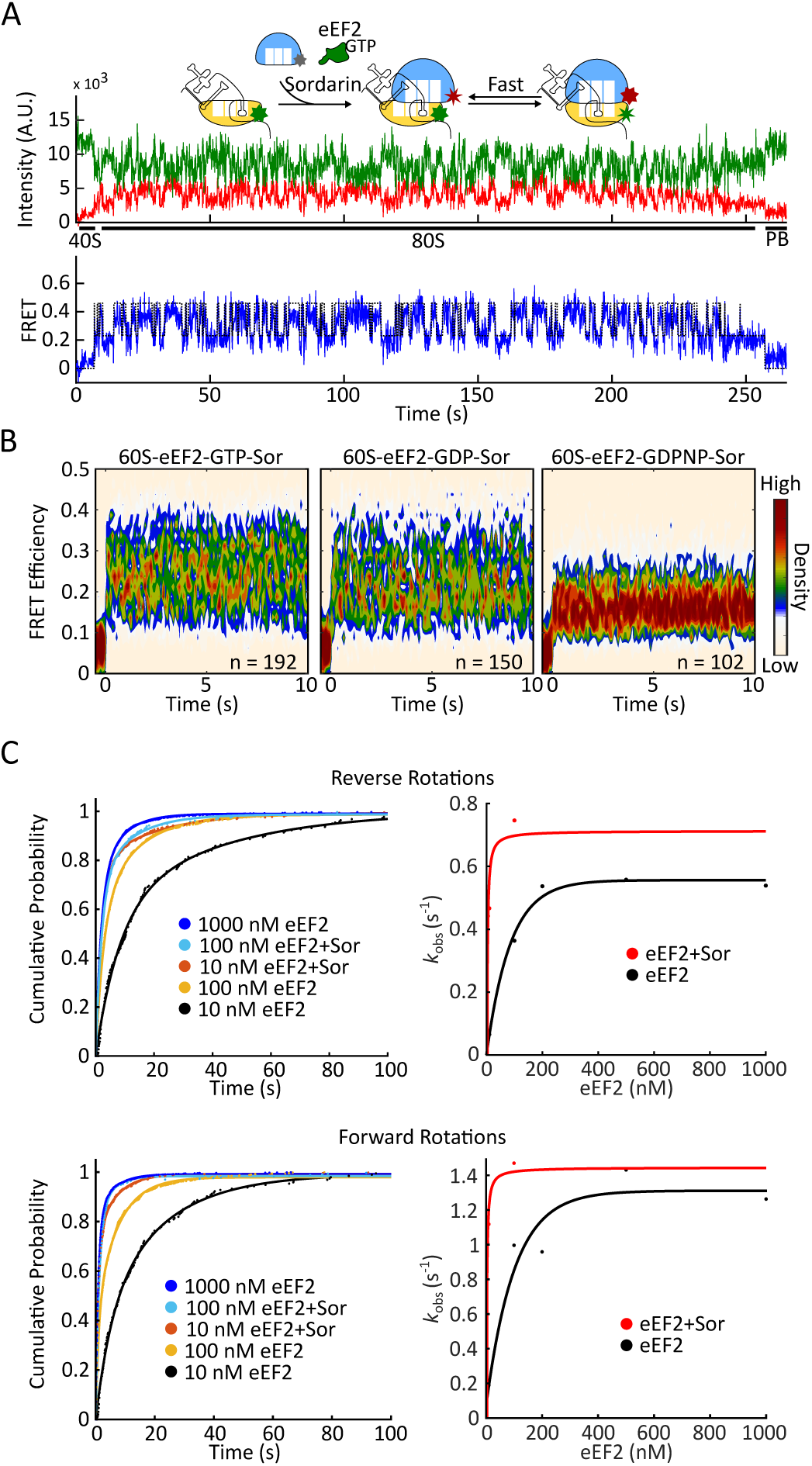
Intersubunit rotations accelerate in the presence of sordarin. A. Cartoon schematic and example experimental trace of 80S-CrPV-eEF2-GTP-Sordarin complex dynamic. 40S-IRES complex were immobilized to the slide and 60S subunits, eEF2, sordarin, and GTP were co-delivered concurrently at the start of observation. Ribosomes continuously oscillated between the non-rotated and rotated states. B. Post-synchronization plot of population FRET efficiency, synchronized to 60S ribosomal subunit arrival (set as zero time point) that rotations were caused by eEF2 in post hydrolysis state. C. Cumulative probability plot shows the comparison of reverse (top) and forward rotation (bottom) dwell times with and without sordarin. Rotations saturated at around 10 nM eEF2 in the presence of sordarin and at 200 nM eEF2 with the drug. Lines represent double-exponential fitting and the resulted rates were fit by hyperbolic functions (red, plus sordarin; black, without sordarin).

The Cryo-EM structure of 80S ribosomes with TSV IRES, eEF2, and sordarin provide us with structural information needed to connect ribosomal rotations to the functional states of the ribosome in the presence of sordarin. As showed in 80S-TSV IRES-eEF2-sordarin complex structure, the degree of subunit rotation correlates with the position of the PKI (Supplementary figure 3). The PKI shows a gradual movement between the A- and the P-sites as the ribosome rotates. From un-translocated IRES, characterized by the rotated conformation, to the fully translocated IRES, characterized by the non-rotated conformation. Which means the subunit rotations of 80S-TSV IRES-eEF2-sordarin complex are correlated with IRES translocation (46). We used this relationship between IRES translocation and subunit rotations in our data interpretation. In our experiments with sordarin and eEF2, a clockwise rotation always corresponds to forward translocation and a counterclockwise rotation corresponds to reverse translocation. Thus, in the presence of sordarin, 80S-CrPV IRES-eEF2 complexes continuously translocate and back translocate.

EF-G, a prokaryotic analog of eFF2, cannot efficiently exchange nucleotides while being ribosome bound (75), suggesting that in the presence of sordarin the energy of GTP hydrolysis is not required for repeated rounds of forward and reverse translocation. To determine if GTP hydrolysis provides energy to IRES translocation, we repeated experiments with GDPNP and GDP. In the presence of sordarin, more than 80% of 80S-IRES-eEF2-GDPNP complexes were characterized by stable FRET (Figure 5A, Supplementary Table 5), which is consistent with the previous proposal that sordarin acts post-GTP hydrolysis (45, 60). On the other hand, eEF2-sordarin promoted subunit rotations in the presence of GDP (Figure 4B, 5A). The first conformational transition in the presence of GDP and sordarin was slightly slower taking 8.4 seconds (vs 6 s in the presence of GTP), while the subsequent transitions were indistinguishable from transitions in the presence of GTP (Figure 5B, Supplementary Figure 6). Thus, in the presence of sordarin, GTP hydrolysis is not required for repeated rounds of forward and reverse translocation.

**Figure 5.**
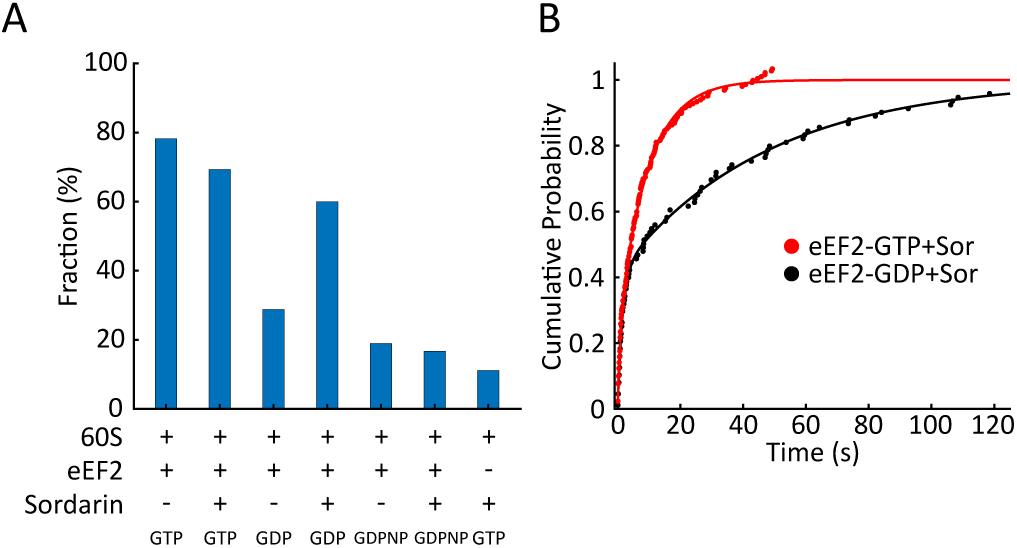
GTP hydrolysis is not required for rotations in the presence of sordarin. A. Fraction of molecules that fluctuated between rotational states (n = 202, 192, 163, 150, 152, 102, and 108, correspondingly). GDPNP suppresses ribosomal rotations in the absence and presence of sordarin consistent with sordarin acting post hydrolysis. Rotations were prevalent with GTP and in presence of sordarin, showing that energy of GTP hydrolysis was not required. Thus, GTP hydrolysis is needed to achieve conformation in which rotations are unlocked, but not needed for subsequent transition. B. The dwell times before the first rotations were fitted by a double exponential function. The first rotation is slower with GDP, consistent with decreased eEF2-GDP affinity to the ribosome.

To exclude the possibility that eEF2 rebinding is driving these events, we conducted wash-off experiments. In these experiments, immobilized 80S-IRES ribosomes were preincubated with eEF2-GTP either in the presence or absence of sordarin and immobilized on the surface of microscope slide. Then ribosomes were imaged to confirm expected behavior. After 30 seconds of imaging, eEF2 and nucleotides were replaced with buffer that does not contain eEF2 and GTP and observation continued. Prewash imaging served as a control and showed that ribosomes underwent rotations identical to ones seen in the experiments described above. After the wash-off, control ribosomes preincubated without sordarin showed stable FRET, with an average of 0.4 rotations per ribosome (in 30 seconds observation window after the wash). However, ribosomes that were preincubated with sordarin continued to fluctuate between high and low FRET states, with an average of 6.1 FRET transitions per ribosome. (Figure 6, Supplementary Table 5). Because wash buffer did not contain eEF2 and GTP, wash precluded the possibility of eEF2 rebinding to the 80S-IRES complexes or nucleotide exchange. Therefore, repeated rounds of translocation are not caused by eEF2 re-binding, but rather promoted by stably bound eEF2 and are thermally driven. This interpretation is also consistent with translocation rates being faster in the presence of sordarin, where reaction essentially becomes unimolecular, and thus expected to be faster than biomolecular translocation in absence of the drug.

**Figure 6.**
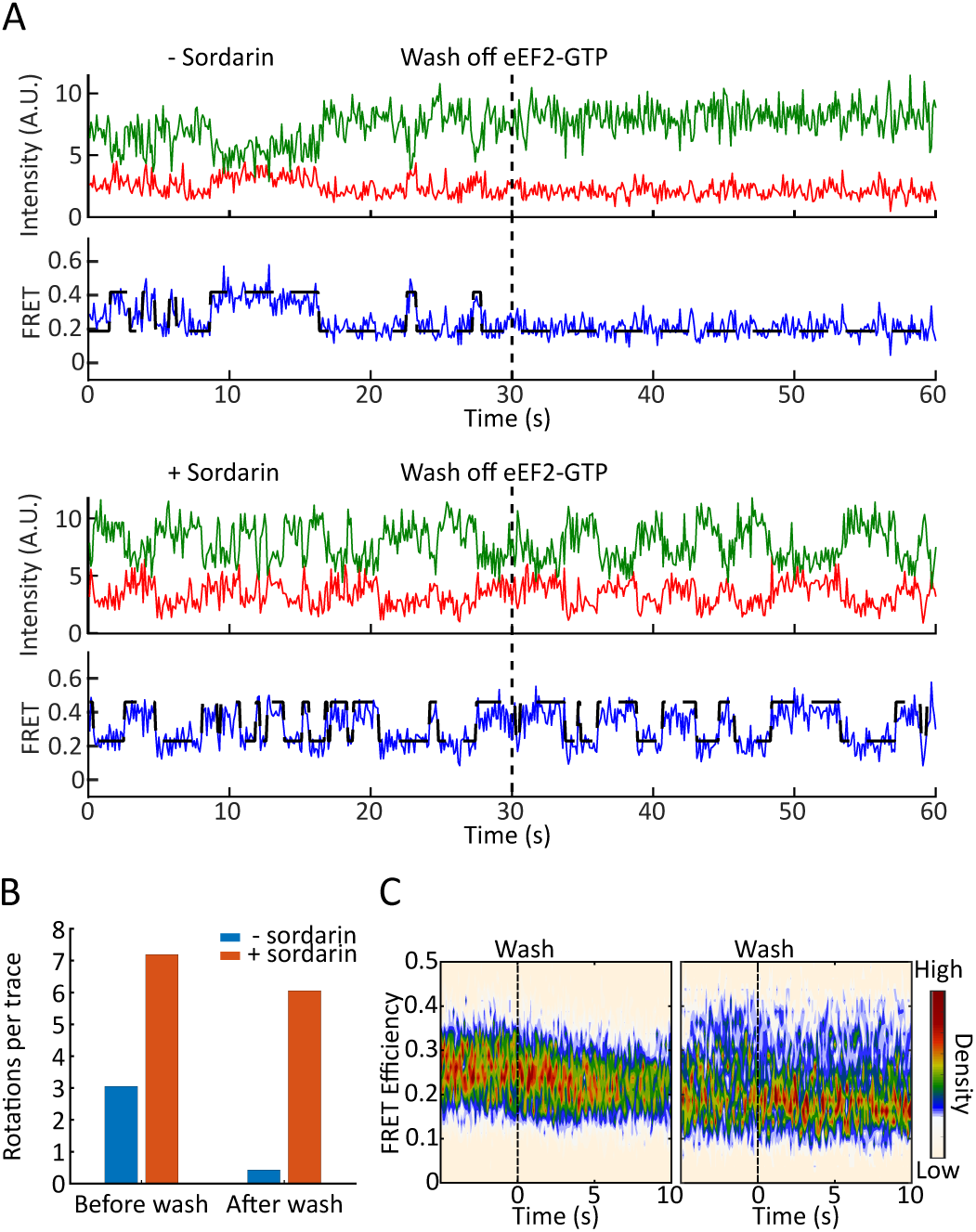
Intersubunit rotations in 80S-CrPV IRES-eEF2 complex are thermally driven in the presence of sordarin. A. Example traces from wash off experiment. 80S-CrPV IRES complex were immobilized to the slide with 100 nM eEF2-GTP without (top) and with (bottom) sordarin (n = 142 and 126, correspondingly). The ribosome oscillates between the non-rotated and rotated states before the wash. After the wash, rotations disappear in the experiment without sordarin, but the ribosome continues fluctuating in the presence of sordarin. B. Number of transitions before and after the wash for eEF2 (blue) and eEF2 plus sordarin (red). C. Traces were synchronized with wash time being set to zero. Ribosomes preincubated with eEF2 (left) rapidly settled in the low FRET state and ribosomes preincubated with eEF2 and sordarin (right) experienced multiple rounds of rotations.

### eEF3 does not affect translocation of CrPV IRES

To investigate the role of yeast elongation factor 3 (eEF3) in IRES translocation, 500 nM eEF3-ATP, Cy5-60S, and 10 or 200 nM eEF2-GTP were co-delivered to immobilized 40S-CrPV complex. Kinetics analysis revealed that the rates of both forward and reverse rotations were unaffected by presence of eEF3 (Supplementary Figure 7). FRET intensity distributions were similarly unaffected. It is possible that eEF3 functions rapidly at timescales faster than the time resolution of our experiments (100 ms). To examine this possibility, we repeated the experiments outlined above in the presence of non-hydrolyzable ATP analog ADPNP. It stabilizes eEF3 on the ribosome and prevents the L1 stalk from opening by eEF3 (43, 76). Similarly, rotation rates and FRET intensity remained unchanged (Supplementary Figure 7). Thus, we concluded that eEF3 has no effects on the translocation of 80S-CrPV IRES complex.

## Discussion

Recent structures of 80S-IRES-complexes mapped structural pathway of translocation, however, the dynamic motions of IRES driven-translation have not been directly observed. Here, we followed intersubunit conformation of eukaryotic ribosomes during initiation and translocation of CrPV IRES in real-time. We assigned the high (0.34) FRET state to the non-rotated ribosomes and the low (0.22) FRET state to the semi-rotated ribosomes, based on the Cryo-EM structures of pre-translocation and post-translocation (18, 48) 80S-CrPV IRES complex. Pre-translocation 80S-IRES complex are conformationally dynamic and spontaneously exchange between the semi-rotated and non-rotated states. This directly confirms a hypothesis, based on Cryo-EM structures, that 80S-CrPV IRES complexes sample different rotational conformations (18, 19). Importantly, 80S-IRES complexes are kinetically heterogeneous as spontaneous exchange was best described by double exponential fits. Furthermore, there were two populations of ribosomes. About 80% of the pre-translocation complexes were predominantly occupying semi-rotated state, while 20% of the pre-translocation complexes mainly occupied a non-rotated state (Supplementary figure 8). It implies that the system is not at a simple two state equilibrium and additional unobserved intermediates might exist.

Spontaneous rotations in the pre-translocation 80S-IRES complex resemble the behavior of tRNA pre-translocation complex, thus indicating similarities between the two types of translocations. The rate of spontaneous rotations in pre-translocation 80S-IRES complex is comparable to that of bacterial pre-translocation ribosomes complexed with tRNA as measured by smFRET studies (40, 77, 78). When eEF2 binds to the 80S-IRES complex, it transiently places ribosome into the rotated conformation. One point of contention is whether eEF2 can bind to the non-rotated conformation or whether eEF2 binding captures the rotated state of the 80S-IRES complex. The ability of translocase to interact with non-rotated ribosomes is well described. EF-G sampling of non-rotated ribosomes with empty A-sites was detected by single-molecule fluorescence (79) and visualized by Cryo-EM (80). L11-tRNA (81, 82) and S6-L9 FRET (78) indicate that EF-G engages both rotated and non-rotated ribosomes. The ensemble measurements of translocation kinetics argue that EF-G can engage both intersubunit conformations (83). Similarly, eEF2 also can recognize non-rotated ribosomes. eEF2-GDPNP causes a single nucleotide toeprint shift in the 5ʹ direction during post-translocation in non-rotated ribosomes (84). At a high concentration of E-site tRNA, eEF2 can promote reverse translocation, a reaction in which a factor is also expected to engage non-rotated ribosomes (85). Our kinetics analysis showed eEF2 promoted both reverse and forward subunit rotations in a concentration dependent manner, which also suggests eEF2 can efficiently recognize both semi-rotated and non-rotated ribosomes. Together, these results demonstrate that the ability of translocase to interact with non-rotated ribosomes is conserved between prokaryotic and eukaryotic elongation as well as in tRNA and IRES translocation.

### eEF3 in IRES translocation

The lack of eEF3 effects on CrPV translocation are in line with our understanding of eEF3 function. During tRNA translocation, the L1 stalk contacts the elbow of the P-site tRNA. Post-translocation eEF3 opens the L1 stalk, thus allowing E-site tRNA dissociation. In the pre-translocation 80S-IRES complex, the bulk of domain I of CrPV IRES displaces the L1 stalk, placing it in a position that is more open than in the tRNA pre-translocation complex (17). The translocated IRES pushes the L1 stalk further open, beyond to what is found in post-translocation ribosomes complexed with tRNA (48). This opening of the L1 stalk is a result of outward movements of domains I and II from the E-site that occur concurrently with reverse head swivel during the late stages of IRES translocation (19). Thus, opening of the L1 stalk is a part of IRES translocation, which explains why eEF3 had no effect on dynamics of intersubunit rotations and, congruently, had no effect on recruitment of the first tRNA (53).

### Role of eEF2 in IRES translocation

In the presence of eEF2 and GTP, 80S-CrPV IRES complexes repeatedly translocated and back translocated. Ribosomes became static after eEF2 was washed off, indicating that repeated translocation events were due to multiple rounds of eEF2-ribosome interactions. Both reverse and forward rotations are promoted by eEF2 and GTP hydrolysis is required (at least for forward translocation), as eEF2-GDP had no effect on ribosome conformation (Supplementary Figure 6A). In prokaryotes, translocation in the presence of GDPNP is slower than in the presence of GTP (75, 79). Similarly, less than 20% 80S-IRES-eEF2 complex underwent translocation in the presence of GDPNP, as most ribosomes were stabilized in the fully rotated state (Figure 3B, 5A). Thus, eukaryotic IRES translocation is remarkably similar to tRNA translocation.

The Brownian mechanism of translocation has been long proposed (86, 87). There the movement occurs stochastically due to Brownian forces, and directionality is provided by recuperating energy. It is supported by a number of studies (reviewed in (88, 89)). Our results show that, ribosome spontaneously translocated and back translocated in the presence of eEF2 and sordarin. Neither reaction required GTP hydrolysis or phosphate release, as shown by wash experiments. The spontaneous and rapid nature of these reactions indicates that sordarin captures the ribosome in the unlocked state, where IRES translocation occurs over a flat energy landscape, as was previously suggested (46). Our results support the Brownian mechanism by directly showing that mid and late stages of translocation are thermally driven in the presence of sordarin.

Remarkably, after the first slower translocation event in the presence of GDP and sordarin, subsequent rounds of translocation were as fast as translocation in the presence of GTP (Supplementary Figure 7). This suggests that GTP hydrolysis is used to achieve the unlocked state. Once the ribosome is unlocked, IRES translocation is thermally driven in the presence of the drug, with subsequent events, such as eEF2 compaction and dissociation possibly providing the recuperating energy.

The CrPV IRES mimics tRNA and translation factors. Despite the ability of the IRES to trigger conformational changes in the ribosome associated with initiation and elongation, the dynamics and conformation of the IRES pre-translocation complex are different from tRNA translocation. Further investigation of the dynamics of eukaryotic tRNA translocation, specifically high-resolution structures of 80S tRNA complex with eEF2 and sordarin, will be needed to see if tRNA translocation is thermally driven by eEF2-sordarin complex.

## ACKNOWLEDGEMENT

We thank Terri Goss Kinzy for providing eEF2- and eEF3-expressing yeast strains. We are grateful to Hong Li, Jinfan Wang and Joseph D. Puglisi for their help and stimulating discussion. We thank Brielle Sorkin for helping with manuscript preparation.

## AUTHOR CONTRIBUTIONS

A.P. and Z.O. designed the study. Z.O. performed all the experiments. A.P. and Z.O. wrote the manuscript.

---

**Supplementary Figure 1.**
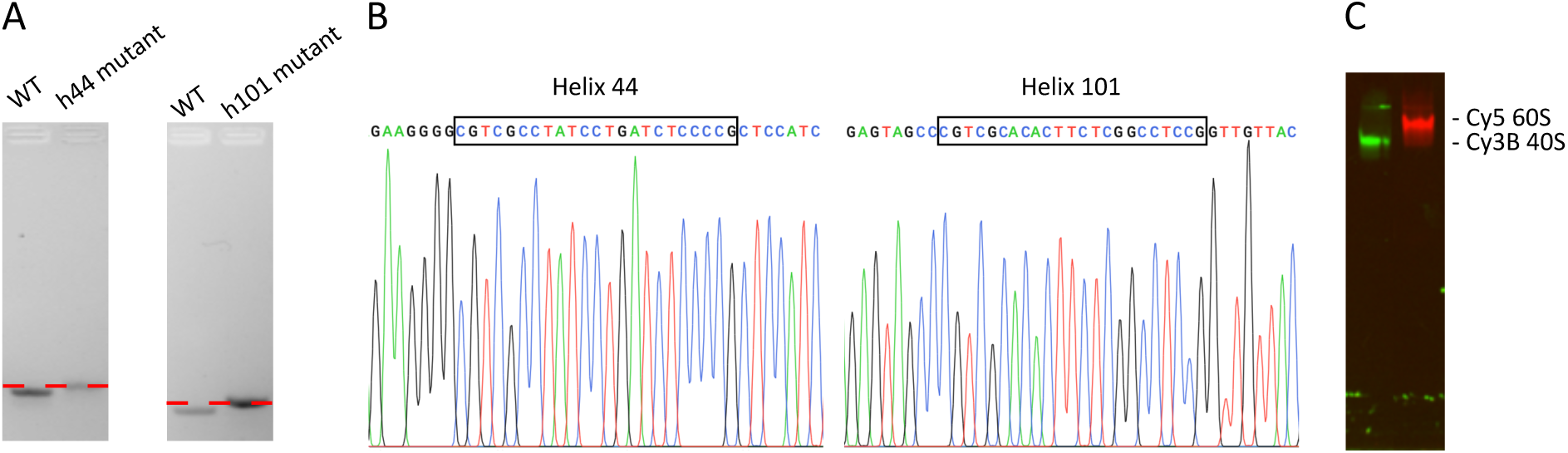
Labeling of the yeast ribosomal subunits. Metastable hairpins were introduced into phylogenetically variable regions of rRNA. The sp68 hairpin was inserted into 18S rRNA, replacing 1699-1702 nucleotides, thus extending h44. The sp22 hairpin was inserted into 25S rRNA, where it replaced nucleotides 3351-3352, thus extending h101. A plasmid-bearing mutated RDNA operon was transformed into yeast and the loss of the wildtype RDN operon carrying plasmid was done, as described in Materials and Methods. The resulting yeast strains that carry the mutated RDNA operon as a sole source of rRNA. A. The presence of the mutated RDN operon was confirmed by PCR with primers flanking either the h44 or h101 regions. The PCR product was ∼19 and 21 nts longer for the mutant hairpins. B. The PCR product was sequenced to further confirm the absence of contamination by the wildtype RDNA. The box indicates the inserted hairpin. C. The labeling and integrity of ribosomal subunits was confirmed by composite agarose-polyacrylamide electrophoresis (47), where purified and labeled ribosomal subunits were migrating as a single band. The labeling efficiency was determined spectrophotometrically and was found to be ∼92% for the 40S subunits and ∼88% for the 60S subunits.

**Supplementary Figure 2.**
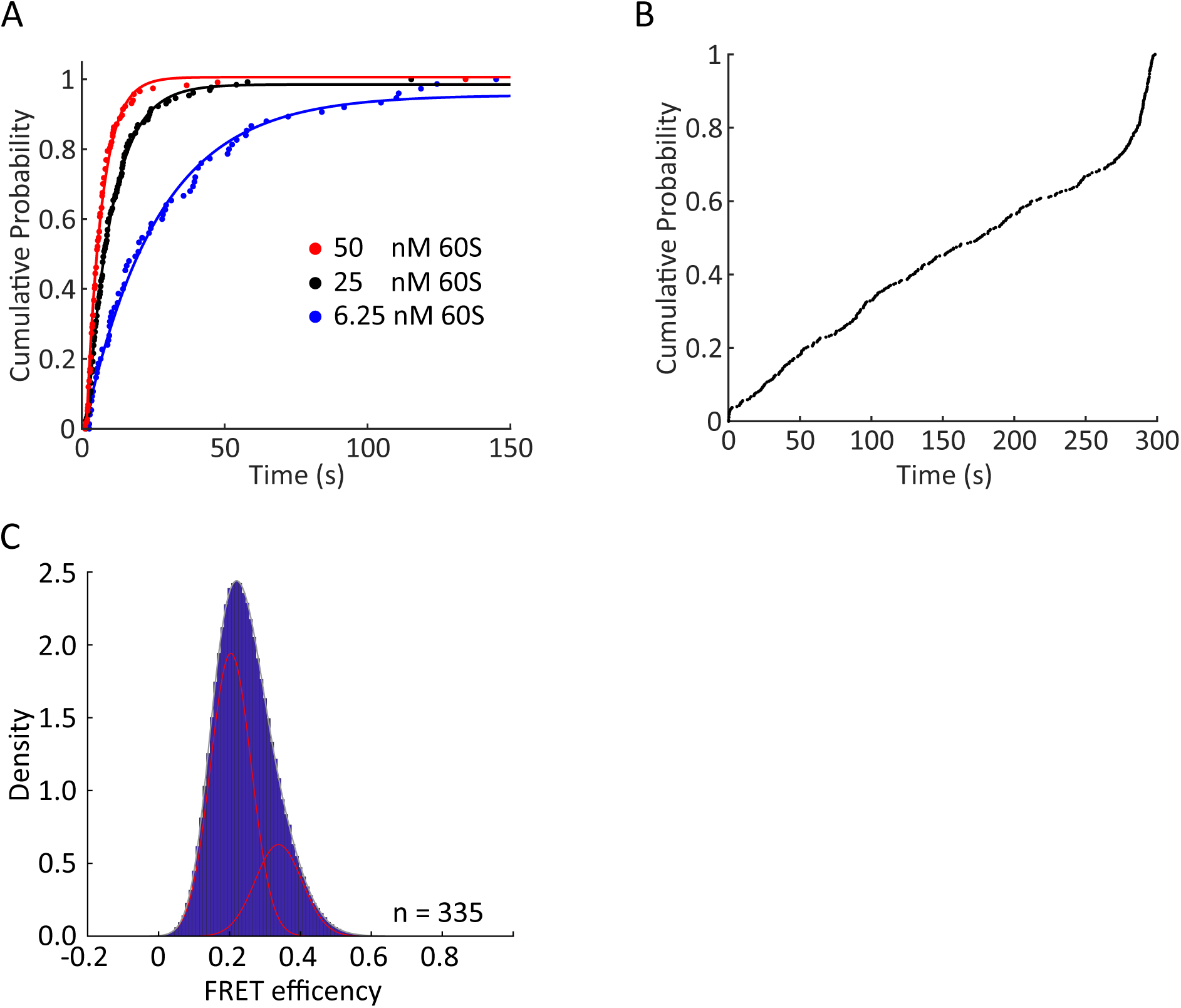
60S subunit arrival to 40S-CrPV IRES complex. A. The cumulative probability of arrival times for 6.25, 25, and 50 nM 60S ribosomal subunits. The empirical Cumulative Distribution Functions were fitted with a single exponential model (*y* = *A*_0_(1 − exp (−*k*_*obs*_ /*t*)+c, n = 102, 130, and 117, correspondingly). B. FRET lifetimes of 60S ribosomal subunit delivery experiment. Hockey stick appearance is due to FRET lasting until the end of the movie (n = 335 molecules). C. The FRET distribution was single modal with a minor right shoulder, indicating that ribosomes prefer a low FRET state. Double Gaussian fit estimated mean FRET efficiency of 0.2 ± 0.06 (95% CI) for low FRET state and 0.33 ± 0.07 (95% CI) for high FRET state correspondingly (n = 335 molecules).

**Supplementary Figure 3.**
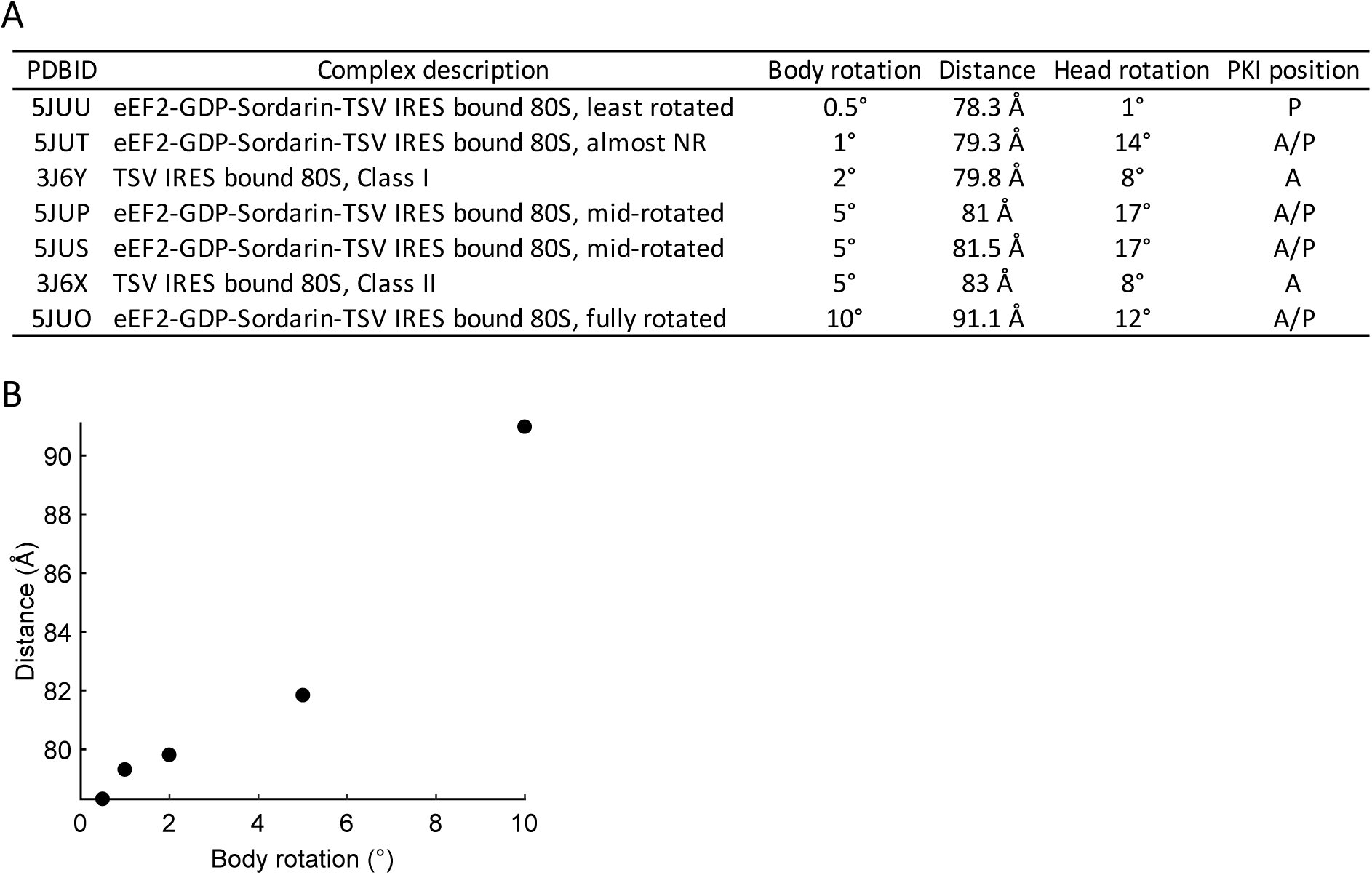
Interpretation on the FRET states. A. Estimated distances between the two labelling sites on helix 44 of 18S rRNA and helix 101 of 25S rRNA were measured as a distance from C4 of the ribose of 1698G to C4 of the ribose of 3350C based on published yeast 80S structures. The structures of helix 44 and helix 101 are not fully resolved in pre-translocation 80S-CrPV IRES structures. So, we used the relevant structure of the pre-translocated 80S-TSV IRES to measure the distance. B. The relationship between intersubunit conformation and distance between h44 and h101.

**Supplementary Figure 4.**
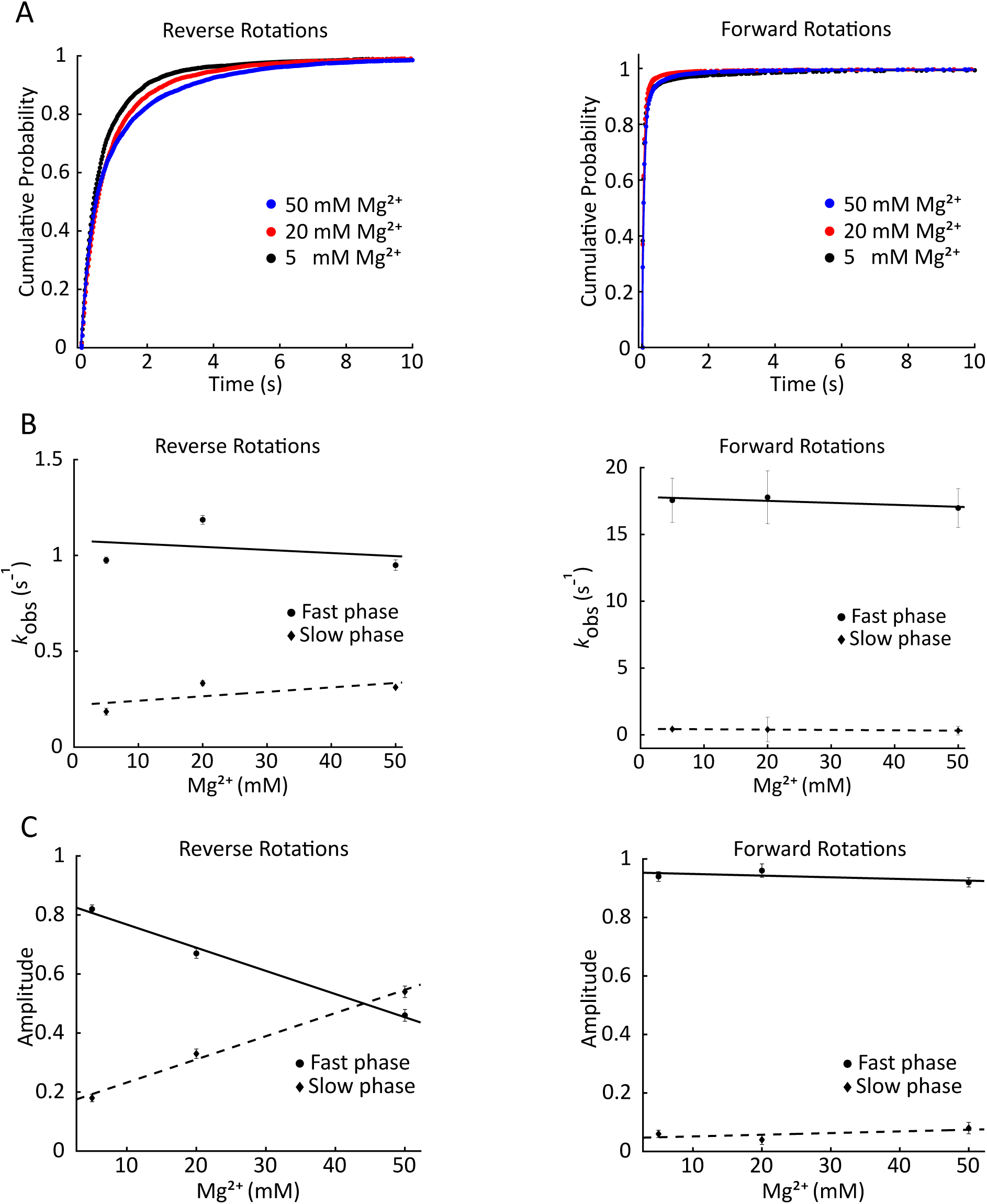
Spontaneous rotations in pre-translocation 80S-CrPV IRES complex become resolved at 25 ms exposure time. A. Cumulative probability plot showed the comparison of reverse (left) and forward rotation (right) dwell times at different magnesium concentrations. Both are best approximated by a double exponential fit. B. The rate for reverse (left) and forward rotations (right) show insignificant changes with the increase of Mg^2+^ concentration. C. Ribosomes prefer the rotated state at elevated Mg^2+^ due to the slow-down of reverse rotation, which is manifested by relative change of amplitudes in the fast and slow phases. The amplitude of double exponential fit for reverse (left) and forward rotations (right) with the change of Mg^2+^ concentration. The amplitude of the fast phase for reverse rotations decreases with the increase of Mg^2+^ concentration (n = 198, 224, and 214 for 5 mM, 20 mM, and 50 mM Mg2+, correspondingly).

**Supplementary Figure 5.**
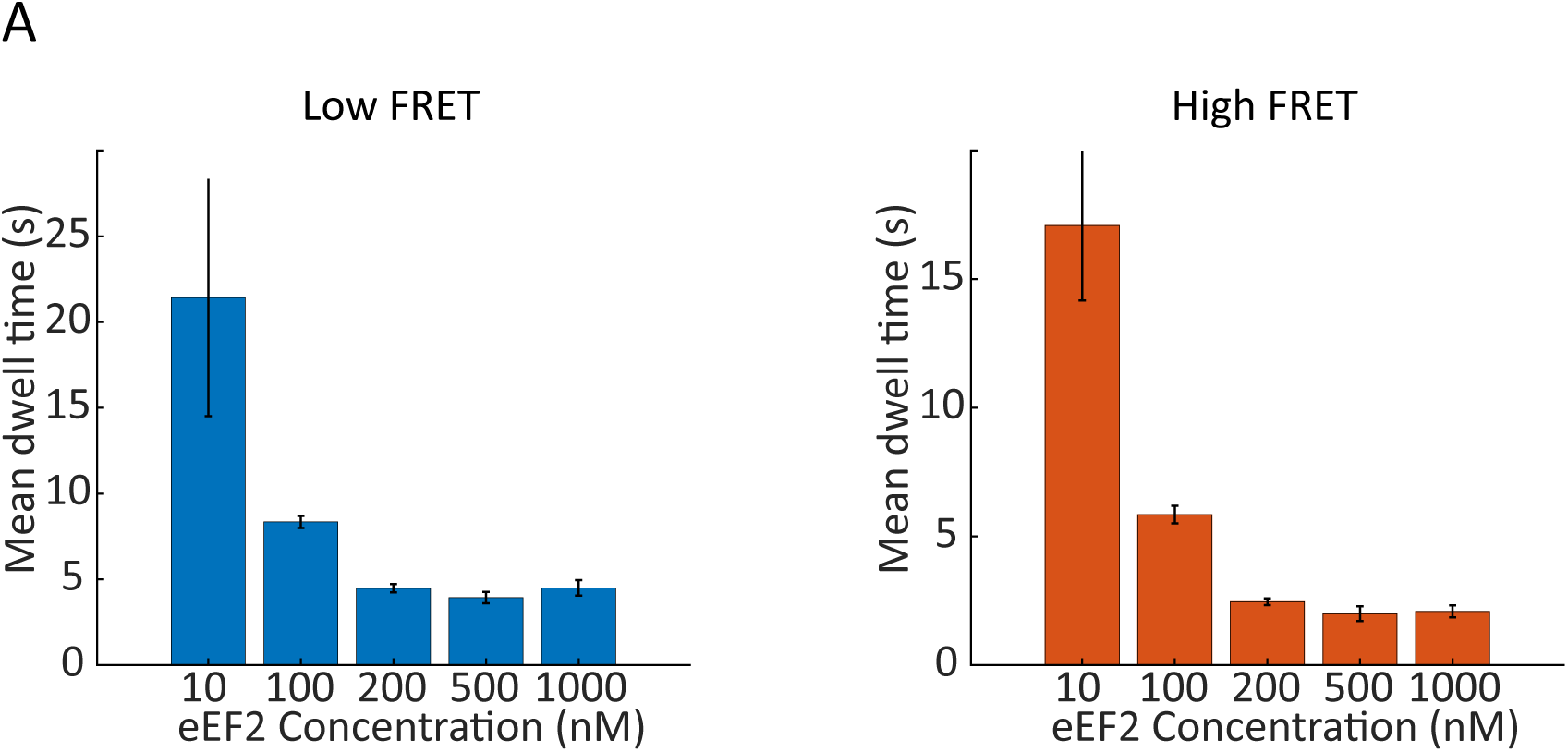
Kinetics of eEF2-induced intersubunit rotations. A. Mean dwell times of the low FRET state (rotated ribosomes, left panel) and the high FRET state (non-rotated ribosomes, right panel) with different eEF2 concentrations. Dwell times decreased with the increase of eEF2 concentration.

**Supplementary Figure 6.**
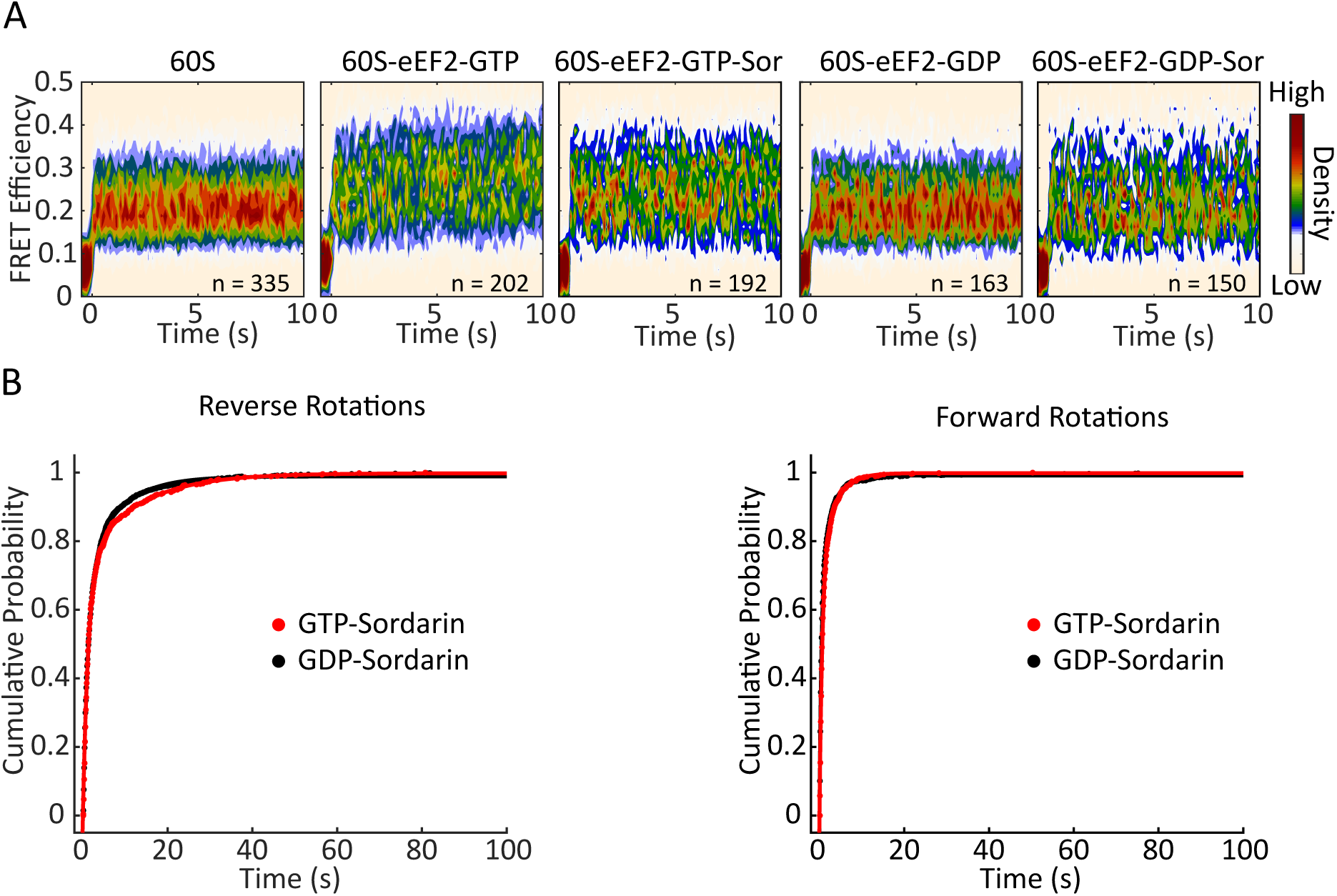
GTP is not required for subunit rotations in the presence of sordarin. A. Post-synchronization plot of FRET efficiency, synchronized to 60S ribosomal subunit arrival (zero time point). eEF2-GDP has no effect on ribosome conformations. B. Cumulative probability plot showed comparison of all subsequent reverse (left) and forward rotation (right) dwell times with the presence of GTP and GDP and have no differences.

**Supplementary Figure 7.**
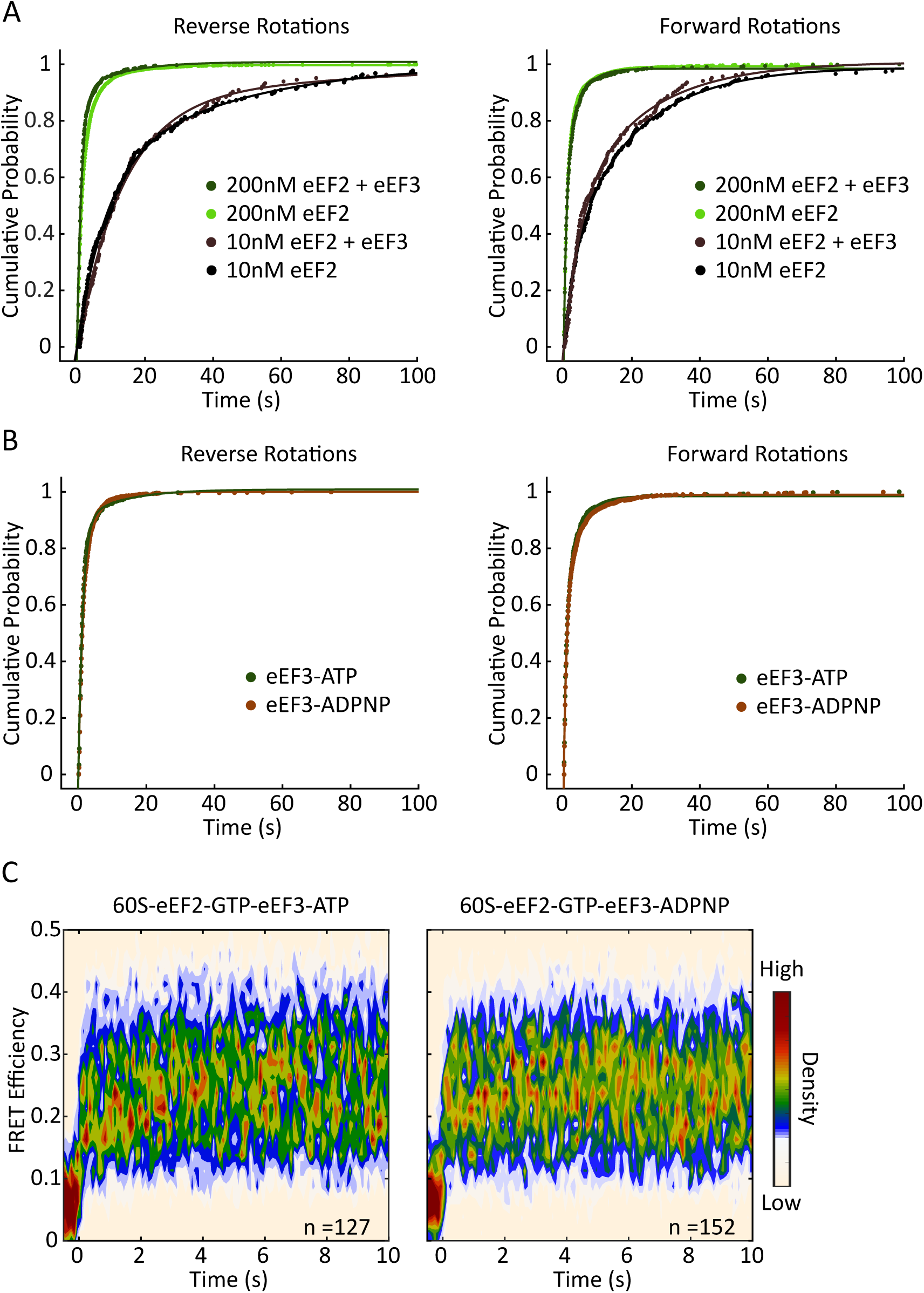
eEF3 does not affect intersubunit dynamics. A. Cumulative probability plot showed comparison of reverse (left) and forward rotation (right) dwell times with and without the presence of eEF3. B. Cumulative probability plot showed comparison of reverse (left) and forward rotation (right) dwell times with the presence of ATP and ADPNP (n = 127 and 152, correspondingly). C. Post-synchronization plot of FRET efficiency observed before and after 60S subunit arrival (n = 127 and 152, correspondingly).

**Supplementary Figure 8.**
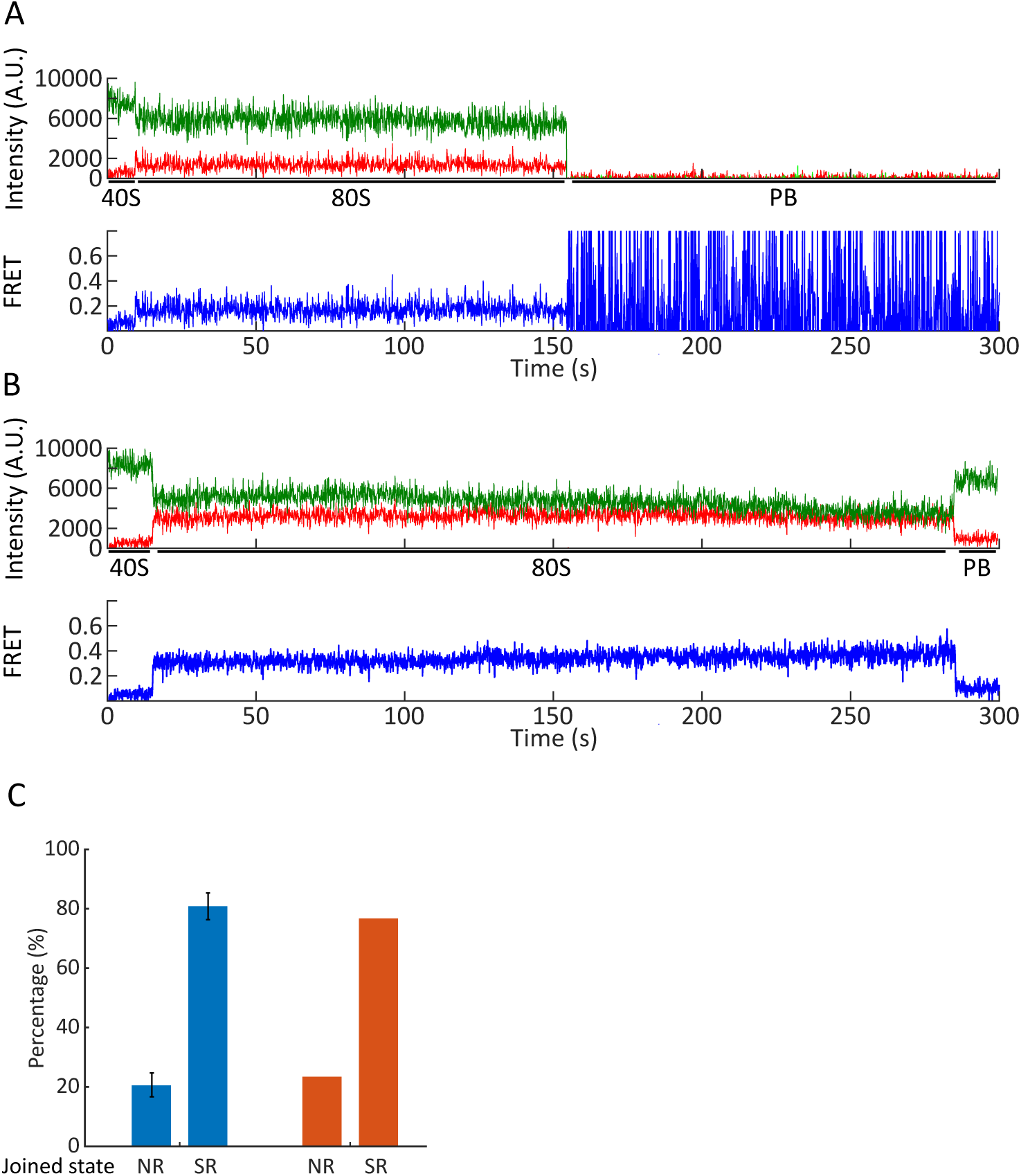
Conformational heterogeneity of pre-translocation 80S-CrPV IRES complex. Example trace for ribosome that joined and remained in semi-rotated state. Spontaneous rotations are masked by camera averaging due to 100 ms exposure. B. Example trace for ribosome that joined and remained in high FRET state. C. Percentage of traces that joined in the non-rotated or rotated state (blue, error bars are 95% CI) and relative fractions from the double Gaussian fit of FRET efficiency (red).

