## Supplemental table for "Sordarin bound eEF2 unlocks spontaneous forward and reverse translocation on CrPV IRES"

| **Name** | **Sequence** | **Restriction site** |
|---|---|---|
| F_h44_SexAI | ACTCACCAGGTCCAGACACAATAA | SexAI |
| R_h44_MluI | GGTGTAAAACCCATACGCGTAATG | MluI |
| F_h101_b | TAGTGGATCCCCCGGGTAACCCAG | BamHI |
| R_h101.3_b | AGCCCGTCGCACACTTCTCGGCCTCCGGTTGTTACGATCTGCTGAGATTAA | |
| F_h101.3_a | CAACCGGAGGCCGAGAAGTGTGCGACGGGCTACTCTACTGCTTACAATACC | |
| R_h101_a | CATACGCGTAATGAAAGTGAACGTAGGTTG | MluI |

**Supplementary Table 2**

Related to Fig. 1C and Supp. Fig. 3A

**60S arrival rates and amplitudes from single-exponential fits**

| Tethered complex (21 °C) | 60S (nM) | Amp | Amp 95% C.I. | Arrival rate (per s) | Rate 95% C.I. | Traces (n) | Median (s) | Mean (s) | Mean 95% C.I. |
|---|---|---|---|---|---|---|---|---|---|
| 40S-CrPV | 6.25 | 0.93 | 0.91 tp 0.95 | 0.034 | 0.031 to 0.036 | 102 | 19.9 | 33.9 | 27.4 to 40.3 |
| 40S-CrPV | 25 | 1.10 | 1.085 to 1.111 | 0.114 | 0.111 to 0.117 | 130 | 7.8 | 12.0 | 9.6 to 14.4 |
| 40S-CrPV | 50 | 1.16 | 1.12 to 1.19 | 0.187 | 0.175 to 0.199 | 117 | 5.3 | 8.3 | 5.8 to 10.7 |

Amp. = Amplitude

Related to Supp. Fig. 4

| Tethered complex (21 °C) | Mg (nM) | Traces (n) | Rotating molecules | Percentage (%) | 95% C.I. | Frame rate | Rotations per trace |
|---|---|---|---|---|---|---|---|
| 40S-CrPV | 5 | 335 | 90 | 26.9 | 0.21 to 0.30 | 100ms | 2.3 |
| 80S-CrPV | 5 | 198 | 164 | 82.8 | 0.77 to 0.88 | 25ms | 23.3 |
| 80S-CrPV | 20 | 224 | 200 | 89.3 | 0.84 to 0.93 | 25ms | 29.5 |
| 80S-CrPV | 50 | 214 | 187 | 87.4 | 0.82 to 0.92 | 25ms | 42.9 |

**Reverse rotations rates and amplitudes from double-exponential fits**

| Tethered complex (21 °C) | Mg (nM) | Amp.1 | Amp.1 95% C.I. | Fast rate (k1, per s) | k1 95% C.I. | Amp.2 | Amp. 2 95% C.I. | Slow rate (k2, per s) | k2 95% C.I. | Traces (n) | Mean (s) | Median (s) | Frame rate |
|---|---|---|---|---|---|---|---|---|---|---|---|---|---|
| 80S-CrPV | 5 | 0.82 | 0.80 to 0.83 | 0.97 | 0.96 to 0.99 | 0.18 | 0.17 to 0.19 | 0.18 | 0.17 to 0.20 | 198 | 0.97 | 0.83 | 25ms |
| 80S-CrPV | 20 | 0.67 | 0.65 to 0.69 | 1.19 | 1.16 to 1.21 | 0.32 | 0.31 to 0.34 | 0.33 | 0.32 to 0.35 | 224 | 1.18 | 0.90 | 25ms |
| 80S-CrPV | 50 | 0.46 | 0.44 to 0.48 | 0.95 | 0.92 to 0.98 | 0.54 | 0.52 to 0.56 | 0.31 | 0.30 to 0.32 | 214 | 1.37 | 1.33 | 25ms |

**Forward rotations rates and amplitudes from double-exponential fits**

| Tethered complex (21 °C) | Mg (nM) | Amp.1 | Amp.1 95% C.I. | Fast rate (k1, per s) | k1 95% C.I. | Amp.2 | Amp. 2 95% C.I. | Slow rate (k2, per s) | k2 95% C.I. | Traces (n) | Mean (s) | Median (s) | Frame rate |
|---|---|---|---|---|---|---|---|---|---|---|---|---|---|
| 80S-CrPV | 5 | 0.83 | 0.81 to 0.85 | 17.56 | 15.9 to 19.22 | 0.16 | 0.14 to 0.17 | 0.30 | 0.22 to 0.38 | 198 | 0.38 | 0.05 | 25ms |
| 80S-CrPV | 20 | 0.97 | 0.95 to 0.99 | 17.78 | 15.79 to 19.77 | 0.03 | 0.003 to 0.05 | 0.15 | 0.13 to 0.18 | 224 | 0.20 | 0.05 | 25ms |
| 80S-CrPV | 50 | 0.96 | 0.94 to 0.97 | 16.98 | 15.54 to 18.42 | 0.04 | 0.004 to 0.08 | 0.32 | 0 to 0.65 | 214 | 0.27 | 0.05 | 25ms |

**Supplementary Table 3**

Related to Fig. 2C,3C  Supp.Fig.6B

**Reverse rotations rates and amplitudes from double-exponential fits**

| Tethered complex (21 °C) | eEF2 (nM) | Amp.1 | Amp.1 95% C.I. | Fast rate (k1, per s) | k1 95% C.I. | Amp.2 | Amp. 2 95% C.I. | Slow rate (k2, per s) | k2 95% C.I. | Traces (n) | Median (s) | Mean | Mean 95% C.I. |
|---|---|---|---|---|---|---|---|---|---|---|---|---|---|
| 40S-CrPV | 1000 | 0.70 | 0.67 to 0.73 | 0.54 | 0.52 to 0.56 | 0.29 | 0.26 to 0.32 | 0.13 | 0.12 to 0.14 | 143 | 1.8 | 4.5 | 2.8 to 6.2 |
| 40S-CrPV | 500 | 0.75 | 0.72 to 0.77 | 0.56 | 0.54 to 0.58 | 0.25 | 0.23 to 0.27 | 0.12 | 0.11 to 0.13 | 129 | 1.6 | 3.9 | 2.5 to 5.4 |
| 40S-CrPV | 200 | 0.75 | 0.73 to 0.76 | 0.54 | 0.52 to 0.55 | 0.25 | 0.23 to 0.26 | 0.102 | 0.095 to 0.109 | 268 | 1.7 | 4.5 | 3.3 to 5.6 |
| 40S-CrPV | 100 | 0.59 | 0.56 to 0.61 | 0.36 | 0.35 to 0.38 | 0.40 | 0.37 to 0.42 | 0.076 | 0.071 to 0.081 | 202 | 3.2 | 8.4 | 7.1 to 9.58 |
| 40S-CrPV | 10 | 0.59 | 0.52 to 0.66 | 0.12 | 0.11 to 0.13 | 0.41 | 0.35 to 0.37 | 0.025 | 0.019 to 0.031 | 169 | 10.4 | 21.4 | 15.6 to 27.3 |

Amp. = Amplitude

**Forward rotations rates and amplitudes from double-exponential fits**

| Tethered complex (21 °C) | eEF2 (nM) | Amp.1 | Amp.1 95% C.I. | Fast rate (k1, per s) | k1 95% C.I. | Amp.2 | Amp. 2 95% C.I. | Slow rate (k2, per s) | k2 95% C.I. | Traces (n) | Median (s) | Mean | Mean 95% C.I. |
|---|---|---|---|---|---|---|---|---|---|---|---|---|---|
| 40S-CrPV | 1000 | 0.80 | 0.77 to 0.82 | 1.26 | 1.22 to 1.31 | 0.20 | 0.18 to 0.22 | 0.20 | 0.17 to 0.22 | 143 | 0.7 | 2.09 | 1.9 to 2.3 |
| 40S-CrPV | 500 | 0.78 | 0.75 to 0.81 | 1.43 | 1.36 to 1.50 | 0.21 | 0.18 to 0.24 | 0.26 | 0.22 to 0.29 | 129 | 1.7 | 2 | 1.7 to 2.3 |
| 40S-CrPV | 200 | 0.79 | 0.77 to 0.80 | 0.96 | 0.94 to 0.98 | 0.21 | 0.19 to 0.22 | 0.17 | 0.16 to 0.18 | 268 | 1 | 2.46 | 2.3 tp 2.6 |
| 40S-CrPV | 100 | 0.43 | 0.42 to 0.45 | 1.00 | 0.94 to 1.05 | 0.55 | 0.53 to 0.56 | 0.133 | 0.129 to 0.138 | 202 | 2.1 | 5.85 | 5.5 to 6.2 |
| 40S-CrPV | 10 | 0.32 | 0.29 to 0.34 | 0.28 | 0.26 to 0.3 | 0.67 | 0.65 to 0.69 | 0.051 | 0.048 to 0.053 | 169 | 7.9 | 17.08 | 14.2 to 20 |

**Supplementary Table 4**

Related to Fig. 3C

**Reverse rotations rates and amplitudes from double-exponential fits**

| Tethered complex (21 °C) | 60S (nM) | eEF2 (nM) | Amp.1 | Amp.1 95% C.I. | Fast rate (k1, per s) | k1 95% C.I. | Amp.2 | Amp. 2 95% C.I. | Slow rate (k2, per s) | k2 95% C.I. | Traces (n) |
|---|---|---|---|---|---|---|---|---|---|---|---|
| 40S-CrPV | 25 | 100 | 0.74 | 0.72 to 0.75 | 0.75 | 0.72 to 0.77 | 0.26 | 0.25 to 0.27 | 0.08 | 0.07 to 0.09 | 192 |
| 40S-CrPV | 25 | 10 | 0.78 | 0.76 to 0.81 | 0.47 | 0.45 to 0.49 | 0.21 | 0.19 to 0.23 | 0.05 | 0.04 to 0.06 | 133 |
| 40S-CrPV | 25 | 5 | 0.74 | 0.71 to 0.76 | 0.42 | 0.40 to 0.45 | 0.20 | 0.17 to 0.22 | 0.10 | 0.09 t0 0.11 | 130 |

Amp. = Amplitude

**Forward rotations rates and amplitudes from double-exponential fits**

| Tethered complex (21 °C) | 60S (nM) | eEF2 (nM) | Amp.1 | Amp.1 95% C.I. | Fast rate (k1, per s) | k1 95% C.I. | Amp.2 | Amp. 2 95% C.I. | Slow rate (k2, per s) | k2 95% C.I. | Traces (n) |
|---|---|---|---|---|---|---|---|---|---|---|---|
| 40S-CrPV | 25 | 100 | 0.72 | 0.65 to 0.78 | 1.47 | 1.32 to 1.62 | 0.28 | 0.21 to 0.35 | 0.30 | 0.22 to 0.38 | 192 |
| 40S-CrPV | 25 | 10 | 0.73 | 0.70 to 0.75 | 1.12 | 1.07 to 1.17 | 0.26 | 0.24 to 0.28 | 0.15 | 0.13 to 0.18 | 133 |
| 40S-CrPV | 25 | 5 | 0.70 | 0.68 to 0.73 | 1.01 | 0.98 to 1.05 | 0.30 | 0.27 to 0.32 | 0.15 | 0.13 to 0.17 | 130 |

**Supplementary Table 5**

Related to Fig. 4A

| Tethered complex (21 °C) | eEF2 (nM) | Traces (n) | Rotating molecules | Percentage (%) | 95% C.I. | Nucleotide | Sordarin |
|---|---|---|---|---|---|---|---|
| 40S-CrPV | 100 | 202 | 158 | 78.2 | 0.72 to 0.84 | GTP | - |
| 40S-CrPV | 100 | 192 | 132 | 68.8 | 0.62 to 0.75 | GTP | + |
| 40S-CrPV | 100 | 163 | 47 | 28.8 | 0.22 to 0.36 | GDP | - |
| 40S-CrPV | 100 | 150 | 90 | 60.0 | 0.52 to 0.68 | GDP | + |
| 40S-CrPV | 100 | 152 | 29 | 19.1 | 0.13 tp 0.26 | GDPNP | - |
| 40S-CrPV | 100 | 102 | 17 | 16.7 | 0.10 to 0.25 | GDPNP | + |
| 40S-CrPV | - | 108 | 12 | 11.1 | 0.06 to 0.19 | GTP | + |
| 40S-CrPV | - | 335 | 90 | 26.9 | 0.21 to 0.30 | GTP | - |

Related to Fig. 4B

**First rotations rates and amplitudes from double-exponential fits**

| Tethered complex (21 °C) | eEF2 (nM) | Nucleotide | Amp.1 | Amp.1 95% C.I. | Fast rate (k1, per s) | k1 95% C.I. | Amp.2 | Amp. 2 95% C.I. | Slow rate (k2, per s) | k2 95% C.I. | n | Sordarin |
|---|---|---|---|---|---|---|---|---|---|---|---|---|
| 40S-CrPV | 100 | GTP | 0.23 | 0.21 to 0.25 | 1.27 | 1.07 to 1.46 | 0.77 | 0.76 to 0.79 | 0.11 | 0.10 to 0.11 | 192 | + |
| 40S-CrPV | 100 | GDP | 0.41 | 0.40 to 0.42 | 0.64 | 0.60 to 0.68 | 0.59 | 0.58 to 0.60 | 0.02 | 0.019 to 0.023 | 163 | + |

Amp. = Amplitude

Related to Fig. 5

**Rotations per trace**

| Tethered complex (21 °C) | Before wash | After wash | n | Sordarin |
|---|---|---|---|---|
| 80S-CrPV-eEF2-GTP | 3.1 | 0.4 | 142 | - |
| 80S-CrPV-eEF2-GTP | 7.2 | 6.1 | 126 | + |
